# Nicotine-mediated recruitment of GABAergic neurons to a dopaminergic phenotype attenuates motor deficits in alpha-synuclein Parkinson’s model

**DOI:** 10.1101/2021.02.15.431290

**Authors:** I-Chi Lai, Benedetto Romoli, Maria Keisler, Fredric P. Manfredsson, Susan B. Powell, Davide Dulcis

**Author notes:** **Corresponding Author:** Davide Dulcis, Department of Psychiatry, UCSD School of Medicine, University of California, San Diego, 9500 Gilman Drive, M/C 0603, Biomedical Science Bldg, La Jolla, CA 92093-0603.

## Abstract

**BACKGROUND:** Previous work revealed an inverse correlation between smoking and Parkinson’s disease (PD) that is associated with nicotine-induced neuroprotection of dopaminergic (DA) neurons against nigrostriatal damage in PD primates and rodent models. Nicotine, a neuroactive component of tobacco, can directly alter the activity of midbrain DA neurons and induce non-DA neurons in the substantia nigra (SN) to acquire a DA phenotype. We investigated the recruitment mechanism of nigrostriatal GABAergic neurons to express DA phenotypes, such as transcription factor Nurr1 and DA-synthesizing enzyme tyrosine hydroxylase (TH), and the concomitant effects on motor function.

**METHODS:** Wild-type and α-syn-overexpressing (PD) mice treated with chronic nicotine were assessed by behavioral pattern monitor (BPM) and immunohistochemistry/*in-situ* hybridization to measure behavior and the translational/transcriptional regulation of neurotransmitter phenotype following selective Nurr1 overexpression or DREADD-mediated chemogenetic activation.

**RESULTS:** Nicotine treatment led to a transcriptional TH and translational Nurr1 upregulation within a pool of SN GABAergic neurons in wild-type animals. In PD mice, nicotine increased Nurr1 expression, reduced the number of α-syn-expressing neurons, and simultaneously rescued motor deficits. Hyperactivation of GABA neurons alone was sufficient to elicit *de novo* translational upregulation of Nurr1 in non-DA neurons. Retrograde labeling revealed that a fraction of these GABAergic neurons projects to the dorsal striatum.

**CONCLUSIONS:** Nicotine exposure initiates neuroprotective mechanisms counteracting the neurodegenerative effects of α-syn accumulation in DA neurons and contributing to Nurr1-mediated therapeutic effects. Revealing the mechanism of nicotine-induced DA plasticity protecting SN neurons against nigrostriatal damage could contribute to developing new strategies for neurotransmitter replacement in PD.

## INTRODUCTION

Parkinson’s disease (PD) is characterized by a progressive neurodegeneration of dopaminergic (DAergic) neurons^1,2^ and aggregation of α-synuclein in the substantia nigra (SN)^3,4^, comprised of the SNc (pars compacta) and the SNr (pars reticulata). Parkinsonism, the collective term for PD motor deficits including bradykinesia, tremor, rigidity, and postural instability, is a consequence of impaired nigrostriatal pathway^5–7^ due to a degeneration of SNc DAergic projection to the dorsal striatum^1,8^. Numerous treatments providing PD symptomatic relief have been developed^9–11^; however, no disease-modifying strategies exist. Neuroprotection targeting classes of compromised neurons in PD and alleviating debilitating movement disorders have been previously investigated^12,13^. Extensive evidence supports an inverse correlation between PD and cigarette smoking^14–17^ and that nicotine mediates neuroprotection when administered before or during nigrostriatal damage both in rodents^18^ and primates^19^.

Nicotine activates nicotinic acetylcholine receptors (nAChRs) and regulates the function of neurons by increasing calcium influx and inducing neuronal depolarization^20,21^. Nicotine-mediated calcium signaling occurs via direct calcium influx through nAChRs, indirect calcium influx through voltage-dependent calcium channels, and intracellular calcium release from internal stores^22–24^. A number of nAChR subtypes are expressed in both DAergic and GABAergic neurons in the SNc and SNr^25–27^ and nicotine-induced neuroprotection can be mediated by heteromeric α4* (primarily α4β2*) and homomeric α7 receptors^28,29^. Importantly, chronic nicotine exposure upregulates α4* nAChRs localized in SNr GABAergic neurons without changing the α4* nAChRs levels in SNc DAergic neurons^26^, suggesting that nicotine might initiate selective activity-dependent signaling on SNc and SNr neurons during chronic exposure.

The expression of the transcription factor Nurr1 (NR4A2), which is essential for the acquisition^30^ and maintenance^31^ of the DAergic phenotype, might participate in the mechanism of nicotine-mediated neuroprotection of nigrostriatal neurons. Studies have shown that Nurr1 expression is regulated by calcium-mediated neuronal activity^32^ and increases in the striatum in response to chronic nicotine administration^33^. Importantly, Nurr1 plays a significant role in neuronal survival^34^ and NR4A-deficient neurons are generally more sensitive to neurodegeneration due to the downregulation of NR4A-dependent neuroprotective gene programs^35^. Emerging evidence indicates that impaired Nurr1 expression might contribute to the pathogenesis of PD^36,37^. Due to its neuroprotective role for DAergic neurons, Nurr1 has been identified as a therapeutic target for PD. Remarkably, it was found that Nurr1 agonists improve behavioral deficits in a PD rat model^38^. Preclinical studies have also shown a promising role of Nurr1 in next-generation PD treatments, including Nurr1-activating compounds and Nurr1 gene therapy aimed at enhancing DA neurotransmission and protecting DAergic neurons from cell damage by environmental toxin and neuroinflammation^37,39,40^.

Here we investigate the cell-specific mechanism through which chronic nicotine influences the activity-dependent regulation of genes controlling the expression of Nurr1 and DA-synthesizing enzyme, tyrosine hydroxylase (TH), in neurons of the SN, while attenuating some of the PD-associated locomotor deficits.

## METHODS

Mice were housed in accordance with the guidelines of the University of California San Diego Institutional Animal Care and Use Committee. An inducible transgenic A53T mutant α-synuclein (hα-syn) mouse PD animal model was utilized to induce pathology in midbrain DAergic neurons. Male and female mice underwent 14 days (P60-to-P74) of chronic nicotine exposure followed by locomotion assessment through the behavioral pattern monitor (BPM). Brain tissue was then processed for either diaminobenzidine (DAB) labelling, fluorescent immunohistochemistry (IHC), or in-situ-hybridization (ISH). Brain injections of retrograde tracer (LumaFluor) and viral vectors to induce Nurr1 overexpression or DREADD-mediated activation were performed in a stereotactic apparatus with a continuous flow of 1% isoflurane. Stereological quantification was performed with Stereologer2000 software. Data were analyzed with SPSS Statistics 26.0 and GraphPad Prism 8.4.0.

Detailed methods are reported in Supplement.

## RESULTS

### Chronic nicotine exposure attenuates PD-associated locomotor deficits and increases Nurr1 expression in the SNr

We used an inducible *Pitx3-IRES2-tTA/tetO-A53T* double transgenic mouse line, which expresses A53T human α-synuclein (hα-syn) in SN DAergic neurons^41^, to investigate whether nicotine exposure improves any behavioral deficit. Breeders were given doxycycline (DOX)-containing (200mg/kg) food pellets, in place of a regular diet, to suppress transgene expression from early embryonic stages through weaning (P21). After weaning (P21), hα-syn+ (positive) mice were placed on normal chow for 90 days (P111) to achieve optimal hα-syn overexpression exclusively in TH+ neurons (Fig. S1A, arrowheads) at P120 (Fig. S1B). At P120, experimental mice began a nicotine consumption (50mg/L nicotine/ 1% saccharin solution) protocol for 14 days (Fig. S2), while control animals were given 1% saccharin solution. Plasma nicotine metabolite levels (18 ng/mL) were assayed by HPLC (NMS Labs). All mice (P125) underwent BPM testing to assess locomotor function. The spatial patterns of locomotion displayed by nicotine-untreated (control) hα-syn+ mice revealed significant differences compared to hα-syn-(negative) mice (Fig. 1), confirming locomotor deficits previously described in this PD mouse model^41^. The hα-syn+ behavioral deficiencies included a number of locomotor and exploratory parameters (Fig. 1A-B), such as distance traveled (two-way ANOVA, hα-syn main effect: F_(1,30)_=8.463, p<0.01), transitions, the number of times mice enter one of nine regions of the testing chamber (two-way ANOVA, hα-syn main effect: F_(1,31)_=8.143, p<0.01), and entries to center (two-way ANOVA, time x hα-syn interaction: F_(3,91)_=3.266, p<0.05, hα-syn main effect: F_(1,32)_=6.114, p<0.05). Remarkably, nicotine-treated hα-syn+ mice did not display these deficits when compared to nicotine-untreated hα-syn+ mice (Fig. 1C, mixed model ANOVA, distance traveled: hα-syn main effect, F_(1,64)_=9.380, p<0.01; transitions: hα-syn x nicotine interaction, F_(1,63)_=5.287, p<0.05, hα-syn main effect, F_(1,63)_=5.841, p<0.05; entries to center: hα-syn x nicotine interaction, F_(1,66)_=4.842, p<0.05). Since the behavioral phenotype in the hα-syn+ mice was stronger in the latter half of the locomotor session, we analyzed 20-40 minutes separately (Fig. 1D, two-way ANOVA, distance traveled: hα-syn main effect, F_(1,65)_=6.930, p<0.05; transitions: hα-syn x nicotine interaction, F_(1,63)_=5.121, p<0.05, hα-syn main effect, F_(1,63)_=6.593, p<0.05; entries to center: hα-syn x nicotine interaction, F_(1,64)_=7.739, p<0.01). The results indicated that chronic nicotine exposure attenuated hα-syn-induced locomotor deficits.

**Figure 1.**
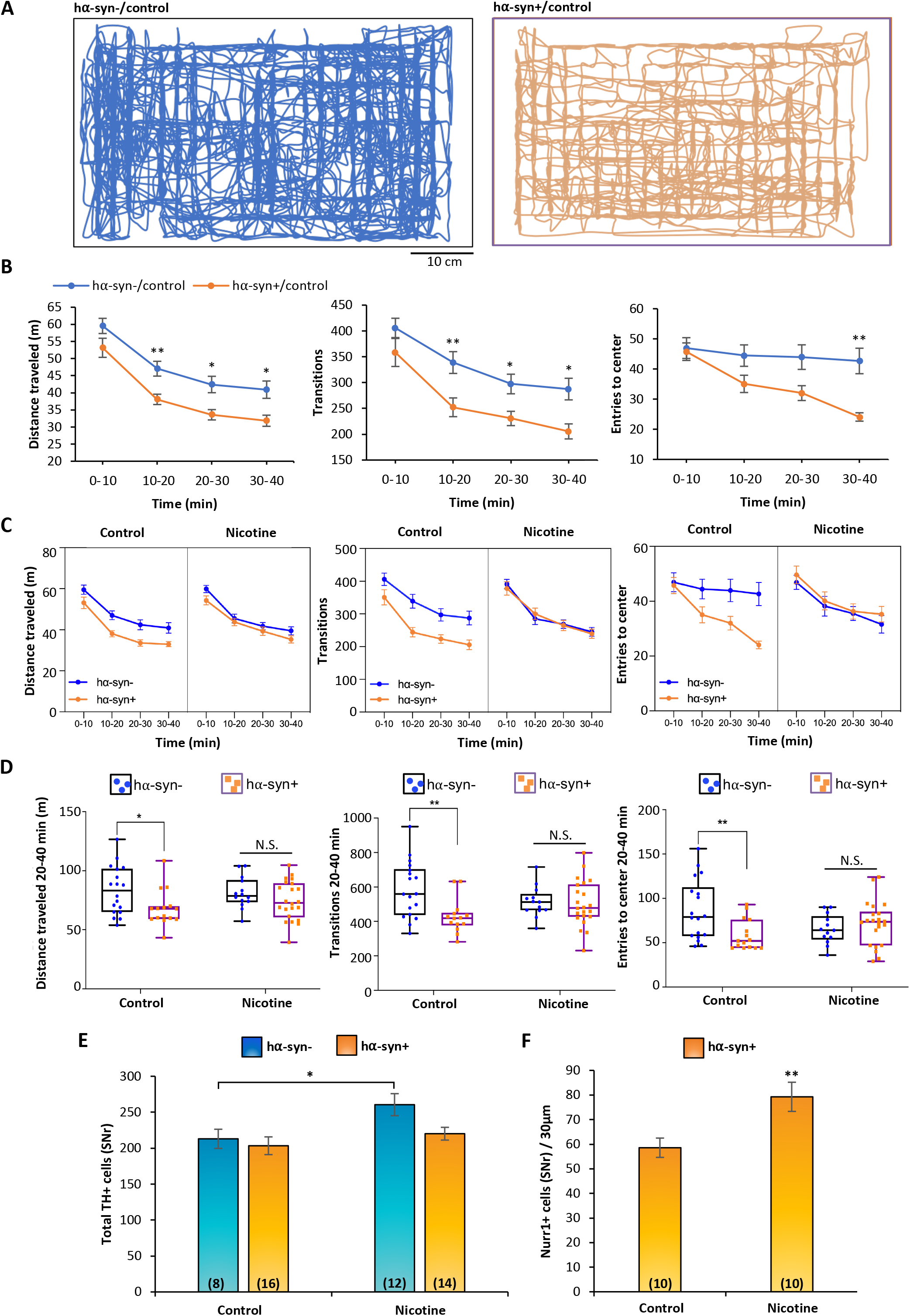
Chronic nicotine exposure attenuates locomotor deficits and increases SNr Nurr1 expression in Pitx3-A53T transgenic mice. **A**, Spatial patterns of locomotion analyzed with the Behavior Pattern Monitor (BPM) within 10-minute duration of the testing session exhibited by a nicotine-untreated (control) Pitx3-A53T transgenic (hα-syn+, right) and a hα-syn-(left) mice. **B**, Locomotion measures from 0 to 40 min of the BPM testing session show that hα-syn+/control mice displayed significant locomotor deficits, including distance traveled (two-way ANOVA, main effect of hα-syn: F_(1,30)_ = 8.463, p<0.01, Bonferroni’s Multiple Comparisons: hα-syn-/control vs hα-syn+/control at 10-20 min: p<0.01, 20-30 min: p<0.05, 30-40: p<0.05), number of transitions across different regions of the chamber (**F**, two-way ANOVA, main effect of hα-syn: F_(1,31)_ = 8.143, p<0.01, Bonferroni’s Multiple Comparisons: hα-syn-/control vs hα-syn+/control at 10-20 min: p<0.01, 20-30 min: p<0.05, 30-40 min: p<0.05), and entries to the center of the chamber (two-way ANOVA, time x hα-syn interaction: F_(3,91)_ = 3.266, p<0.05, main effect of hα-syn: F_(1,32)_ = 6.114, p<0.05, Bonferroni’s Multiple Comparisons: hα-syn-/control vs hα-syn+/control at 30-40 min: p<0.01). Every measure shows a main effect of time, p<0.0001. Graphs show mean ± SEM: *p<0.05, **p<0.01. **C-D**, Chronic nicotine exposure attenuated locomotor deficits as no significant differences in these measures were observed between nicotine-exposed hα-syn- and hα-syn+ groups. **C**, mixed model ANOVA analyzing the effects of hα-syn, nicotine across the 40-minute session on distance traveled: main effect of hα-syn, F_(1,64)_ = 9.380, p<0.01; transitions: hα-syn x nicotine interaction, F_(1,63)_ = 5.287, p<0.05, main effect of hα-syn: F_(1,63)_ = 5.841, p<0.05; entries to center: hα-syn x nicotine interaction, F_(1,66)_ = 4.842, p<0.05. Every measure shows a main effect of time, p<0.0001. **D**, two-way ANOVA performed on the 20-to-40 min interval, distance traveled: main effect of hα-syn, F_(1,65)_ = 6.930, p<0.05, Bonferroni’s Multiple Comparisons: hα-syn-/control vs hα-syn+/control, p<0.05; transitions: hα-syn x nicotine interaction, F_(1,63)_ = 5.121, p<0.05, main effect of hα-syn, F_(1,63)_ = 6.593, p<0.05, Bonferroni’s Multiple Comparisons: hα-syn-/control vs hα-syn+/control p<0.01; entries to center: hα-syn x nicotine interaction, F_(1,64)_ = 7.739, p<0.01, Bonferroni’s Multiple Comparisons: hα-syn-/control vs hα-syn+/control p<0.01. Graphs show all data points with medians and interquartile range. *p<0.05, **p<0.01, N.S., not significant. The number of animals (males and females) for each group is: hα-syn-/control (N=18), hα-syn-/nicotine (N=14), hα-syn+/control (N=17), and hα-syn+/nicotine (N=22). **E**, Stereological quantification revealed that chronic nicotine exposure increased the number of TH+ neurons in hα-syn- but not hα-syn+ mice (t_(18)_=2.18, p<0.05). Graph shows mean ± SEM: *p<0.05. The number of animals is annotated in parenthesis for each condition. **F**, Chronic nicotine exposure increased the number of Nurr1+ cells in the SNr of hα-syn+ mice (t_(18)_=2.91, p<0.01). Graph shows mean ± SEM: **p<0.01. The number of animals is 10 mice/condition.

Because nicotine directly activates SN DAergic neurons via presynaptic nAChRs^42^ and has been shown to exert neuroprotective effects against PD nigrostriatal damage of DA neurons in rodents^18^, we investigated the effect of 2-week nicotine exposure on the number of SN neurons expressing the DA-synthesizing enzyme, tyrosine hydroxylase (TH). As previously found in wild-type mice^43^, unbiased stereological quantification of TH+ neurons in hα-syn- mice revealed that chronic nicotine exposure increased the number of TH+ neurons (Fig. 1E) in the SNr (mean±SEM: control = 213±13, nicotine = 261±15, t_(18)_=2.18, p<0.05) while the SNc was unaffected (mean±SEM: control = 1235±58, nicotine = 1185±63). Importantly, hα-syn+ mice did not respond to nicotine in the same way (Fig. 1E, mean±SEM: control = 203±12, nicotine = 220±9). While SNr TH expression remained unchanged, nicotine-treated hα-syn+ mice showed a higher number of SNr Nurr1+ cells than control hα-syn+ mice (Fig. 1F, mean±SEM: control = 59±4, nicotine = 79±6, t_(18)_=2.91, p<0.01), indicating that a reserve pool^44^ of TH-negative neurons in the SNr acquired the DAergic marker, Nurr1, in response to nicotine exposure.

To determine whether the nicotine-mediated increase in the number of TH-expressing neurons occurs through recruitment of pre-existing SNr neurons to such DAergic phenotype, we tested the effects of 2-week nicotine exposure on the SN of adult wildtype mice (P60). After nicotine exposure, brain tissue was labelled with the DAergic TH, neuronal NeuN, and nuclear DRAQ5 IHC markers. As observed in *Pitx3-IRES2-tTA/tetO-A53T* transgenic mice (Fig.1E), stereological quantification indicated that chronic nicotine exposure in wild-type mice significantly increased the number of TH+ cells (Fig. 2A, arrows, inset) in the SNr (mean±SEM: control = 101±5, nicotine = 148±7, t_(19)_=5.01, p<0.0001, Fig. 2C), but not in the SNc (Fig. 2B). All SNr TH-expressing cells in nicotine-treated mice expressed Nurr1 (Fig. 2D, inset) and NeuN (Fig. 2E, inset) markers. The increased number of TH+ neurons was not due to an increase of neuroproliferation or cell migration, as no change in the total number of DRAQ5+ (mean±SEM: control = 46±5, nicotine = 42±5) and NeuN+ cells (mean±SEM: control = 16±1, nicotine = 17±1) was observed in the nicotine-exposed group (Fig. 2F-G).

**Figure 2.**
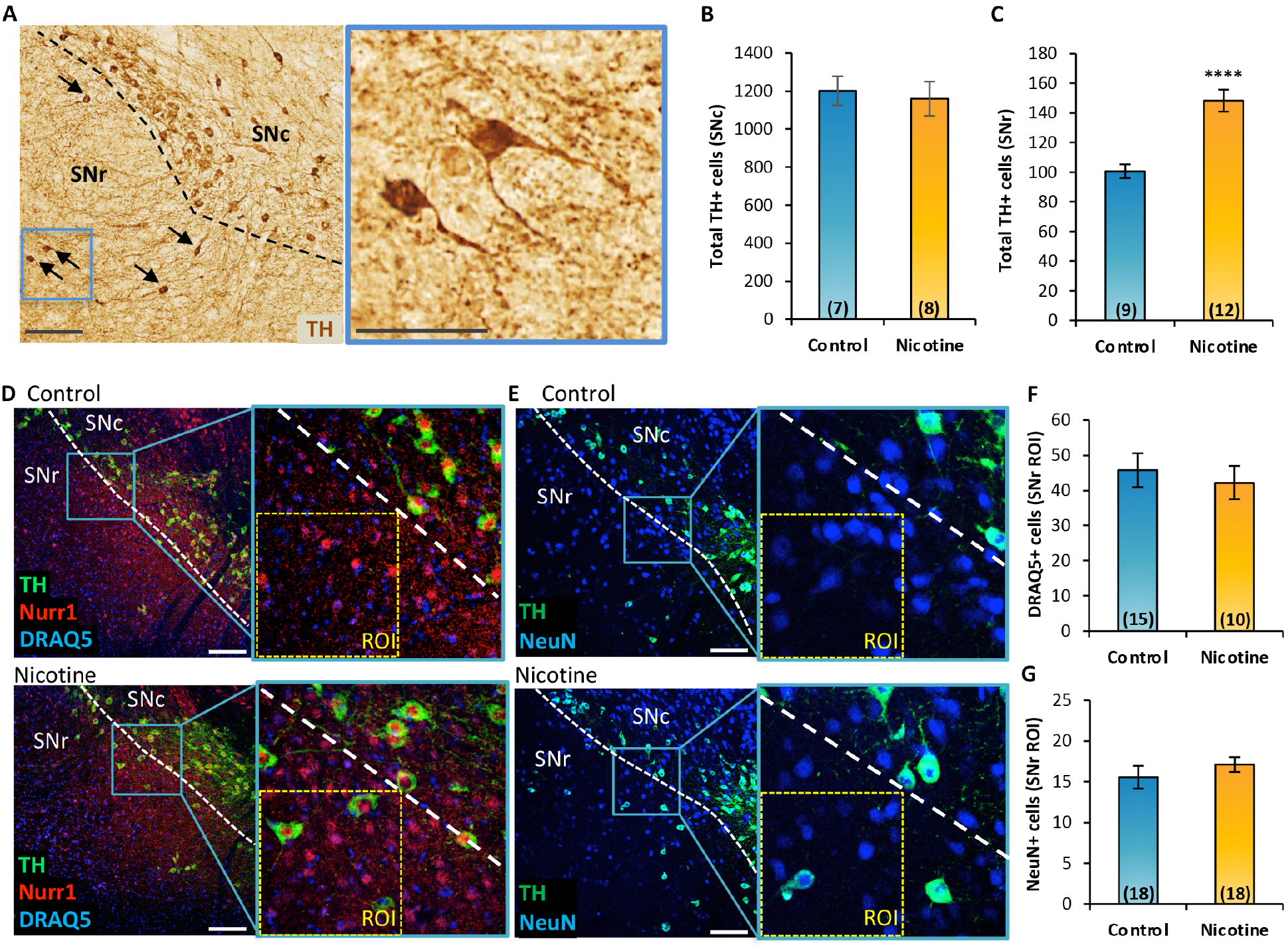
Chronic nicotine exposure increases the number of TH+ cells in the SNr without affecting the number of DRAQ5+ and NeuN+ cells. **A**, Representative SN section of wild-type mice displaying DAB immunoreactivity for TH after chronic nicotine exposure. Arrows indicate TH+ neurons in the SNr. Scale bars = 150 μm; inset, 75 μm. **B-C**, DAB stereological quantification showed that chronic nicotine exposure did not change the number of TH+ cells in the SNc (**B**) but increased the number of TH+ cells in the SNr (**C**, t_(19)_=5.01, p<0.0001). Graphs show mean ± SEM: ****p<0.0001. The number of animals is annotated in parenthesis for each condition. **D-E**, Confocal images showing TH, Nurr1, DRAQ5 (**D**), and NeuN (**E**) immunofluorescence in the SN of control and nicotine-exposed mice. **F-G**, Quantification (SNr ROI) of IHC preparations shown in **D-E** revealed no change in the numbers of DRAQ5+ (**F**) and NeuN+ (**G**) cells. Graph shows mean ± SEM. The number of animals is annotated in parenthesis for each condition. Scale bars = 150 μm. ROI = 150 μm x 150 μm. Abbreviations: ROI, Region of Interest; SNc and SNr, substantia nigra compacta and reticulata.

### Nicotine-induced neurotransmitter plasticity occurs via translational induction of Nurr1 and transcriptional TH regulation in non-DAergic neurons

To understand the level of gene regulation underlying the acquisition of the DAergic phenotype by identified non-DAergic SNr neurons in response to chronic nicotine exposure, we first monitored changes in protein expression of the DA-synthesizing enzyme TH, the glutamate-decarboxylase-67 (GAD67) labeling GABAergic cells, and the transcription factor Nurr1 across conditions (Fig. 3A-B). Nurr1 protein, which is essential for the acquisition and maintenance of the DAergic phenotype^30,31^, was detected in TH+ neurons as expected. However, it was also clearly expressed in TH-negative GABAergic SNr cells, as shown by the GAD67+/Nurr1+ colocalization (Fig. 3A-B, arrowheads). Co-activator Foxa2, which interacts with Nurr1 to promote the survival of midbrain DAergic neurons against toxic insults^40^, displayed overlapping immunoreactivity in Nurr1+/TH-negative cells (Fig. 3C-D, arrowheads). Nicotine-exposed mice (P60) displayed a significant increase in the number of Nurr1+/TH+ cells (Fig. 3B, arrows; Fig. 3E, mean±SEM: control = 100±4 %, nicotine = 139±6 %, t_(21)_=5.51, p<0.0001,) and a concomitant surge in the number of TH-negative neurons co-expressing Nurr1/Foxa2, when compared to controls (mean±SEM: control = 100±5 %, nicotine = 139±10 %, t_(9)_=3.50, p<0.01, Fig. 3D, arrowheads; 3E). While the increase of Nurr1/TH co-expression was expected given the nicotine-mediated increase in TH phenotype (Figs. 1E, 2C), the upregulation of Nurr1/Foxa2 co-expression in non-DAergic neurons could represent part of the priming mechanism generating a molecular memory of nicotine exposure aimed at expanding the reserve pool of potential neurons equipped to undergo the TH genetic program when properly motivated by a persistent nicotine exposure.

**Figure 3.**
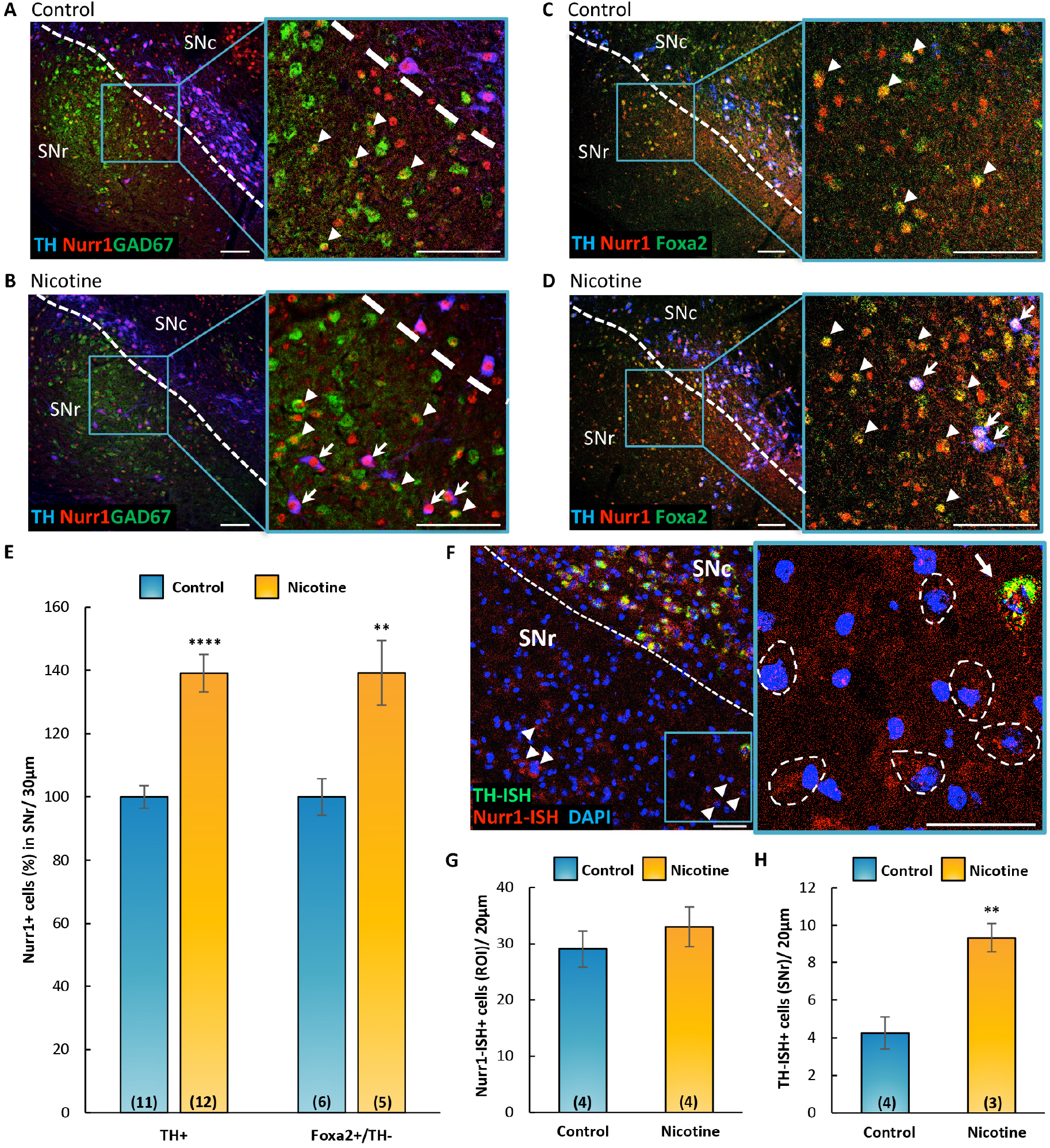
Chronic nicotine exposure increases Nurr1 expression via translational upregulation and induces *de novo* transcription of TH in non-DAergic cells. **A-D**, Confocal images of coronal sections (30 μm) through the SN of control (**A, C**) and nicotine-exposed (**B, D**) mice, labeled with TH, Nurr1, GAD67-GFP, and Foxa2 markers. Insets display immunoreactive TH-/Nurr1+/GAD67+ (**A-B**, arrowheads), TH-/Nurr1+/Foxa2+ (**C-D,** arrowheads), and TH+/Nurr1+ (**B, D,** arrows) cells in the SNr. Scale bars = 100 μm. **E**, Quantification (%) of IHC preparations shown in A-D indicated that chronic nicotine exposure in adult (P60) mice increased the number of TH+/Nurr1+ (t_(21)_=5.51, p<0.0001) and TH-/Foxa2+/Nurr1+ (t_(9)_=3.50, p<0.01) cells in the SNr. Graph shows mean ± SEM: **p<0.01, ****p<0.0001. The number of animals is annotated in parenthesis for each condition. **F**, Confocal image of Nurr1/TH *in situ* hybridization (ISH) of a representative coronal section (20 μm) through the SN of nicotine-exposed adult (P150) mice. DAPI was used to label nuclei. Non-DAergic (TH-ISH negative) Nurr1-ISH+ cells (arrowheads, dashed contours in inset) and TH-ISH+/Nurr1-ISH+ cell (arrow) are observed in the SNr. Scale bars = 50 μm. **G-H**, Quantification of IHC preparations shown in F (ROI, 200 μm x 200 μm) revealed no difference in the number of Nurr1-ISH+ cells between control and nicotine-exposed groups (**G**). Chronic nicotine exposure increased the number of TH-ISH+ cells (**H**, t_(5)_=4.06, p<0.01). Graphs show mean ± SEM: **p<0.01. The number of animals is annotated in parenthesis for each condition. Abbreviations: ROI, Region of Interest; SNc and SNr, substantia nigra compacta and reticulata.

We performed RNAscope *in situ* hybridization (ISH; Fig. 3F) to investigate whether the increased level of Nurr1 and TH protein expression took place via transcriptional or translational regulation. We found that Nurr1 transcripts were present in a significant fraction of TH-ISH-negative neurons (Fig. 3F, arrowheads; inset, dashed contours) in addition to TH-ISH+ neurons (Fig. 3F inset, arrow). This finding was confirmed by quantification data showing no difference in the total number of Nurr1-ISH+ cells across groups (Fig. 3G, mean±SEM: control = 29±3, nicotine = 33±4,), indicating that the increased number of Nurr1+ neurons observed in response to chronic nicotine exposure resulted from translational upregulation. In contrast, the increased number of TH-ISH+ cells in nicotine-exposed mice (Fig. 3H, mean±SEM: control = 4.25±0.85, nicotine = 9.33±0.88, t_(5)_=4.06, p<0.01) revealed *de novo* transcription of TH mRNA in pre-existing SNr non-DAergic neurons.

### The pool of neurons recruitable for nicotine-induced TH plasticity is GABAergic

Because the SNr is primarily composed of GABAergic cells^7,45^, we utilized the vesicular GABA transporter (VGAT)-ZsGreen transgenic mice, which constitutively expresses the ZsGreen-fluorescent protein in all GABAergic cells, to determine the fraction expressing TH and Nurr1 in the control condition. Quantification of TH/VGAT colocalization revealed a coexpression of 27±4% (mean±SEM) in the SNc and 47±4% (mean±SEM) in the SNr (Fig. 4A, inset arrows; Fig. 4B; Video S1) in control conditions. To identify SNr neurons recruitable by nicotine exposure to a TH phenotype, we investigated the pool of VGAT-expressing neurons that co-localized with Nurr1 and NeuN (Fig. 4C). We found that the fraction of Nurr1+/VGAT+/NeuN+ neurons (Fig. 4C, arrowheads) represents 39±2% (mean±SEM) of all SNr VGAT+ neurons (Fig. 4D). This pool of SNr GABAergic neurons, which display the molecular marker Nurr1 even before nicotine exposure, could represent a readily available reserve pool for nicotine-induced TH acquisition and for targeted activity-dependent manipulations.

**Figure 4.**
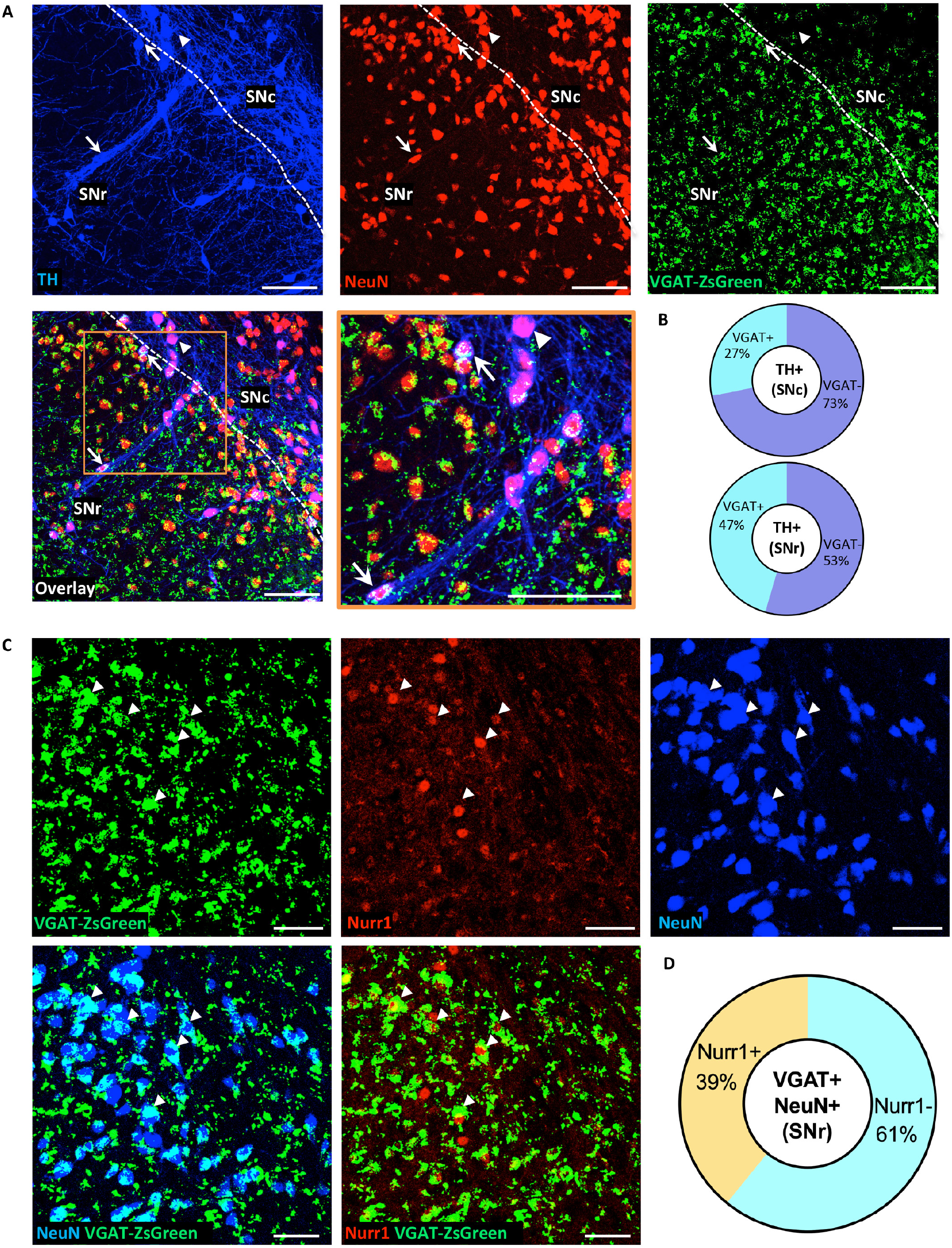
GABAergic neurons expressing Nurr1 in the SNr revealed a reserve pool recruitable to acquire a DAergic phenotype. **A**, Representative confocal images of the SN of adult (P60) vesicular GABA transporter (VGAT)-ZsGreen mice display the distribution of TH+/VGAT+ (arrow) and TH+/VGAT-(arrowhead) neurons. Scale bars = 100 μm. **B**, Quantification of IHC preparations shown in A indicated that 26.7 ± 3.9 % of SNc TH+ neurons and 46.8 ± 4.3 % of SNr TH+ neurons express VGAT. **C**, Confocal images of representative SNr sections show VGAT+/Nurr1+/NeuN+ colocalization (arrowheads). Scale bars = 100 μm. **D**, Quantification of IHC preparations shown in C indicated that 38.8 ± 2.2 % of SNr VGAT+ neurons express Nurr1.

### A fraction of SNr GABAergic Nurr1+ neurons project to the striatum

The nigrostriatal pathway, which is affected by neurodegeneration in PD, comprises DAergic neurons originating from the SNc and projecting to neurons located in striatum subnuclei. However, an additional fraction of nigrostriatal projections originates from GABAergic neurons located in the SNr^46–48^. To confirm the connectivity of GABAergic SNr-to-striatum-projecting neurons, fluorescent retrobeads (555 nm) were injected into the dorsal striatum (Fig. 5A) for a 10-day retrograde tracing of striatal neuronal terminals to their SN somata. Retrobead-labelled cell bodies localized in the SNr identified SNr-to-striatum-projecting neurons (Fig. 5B). Retrobead accumulation was detected in both VGAT+/TH-(Fig. 5B, arrows) and TH+ SNr somata (Fig. 5B inset, arrowhead), in addition to all TH+ SNc-to-striatum-projecting neurons. The schematic diagram (Fig. 5C), generated by compiling our retrograde tracing with IHC data, provides a qualitative representation of the distribution of SNc DAergic and SNr GABAergic neurons and their axonal projections. The fraction of Nurr1+ GABAergic neurons projecting to the striatum could serve as a reserve neuronal pool with the potential to acquire the TH phenotype and in turn replenish DA function in PD.

**Figure 5.**
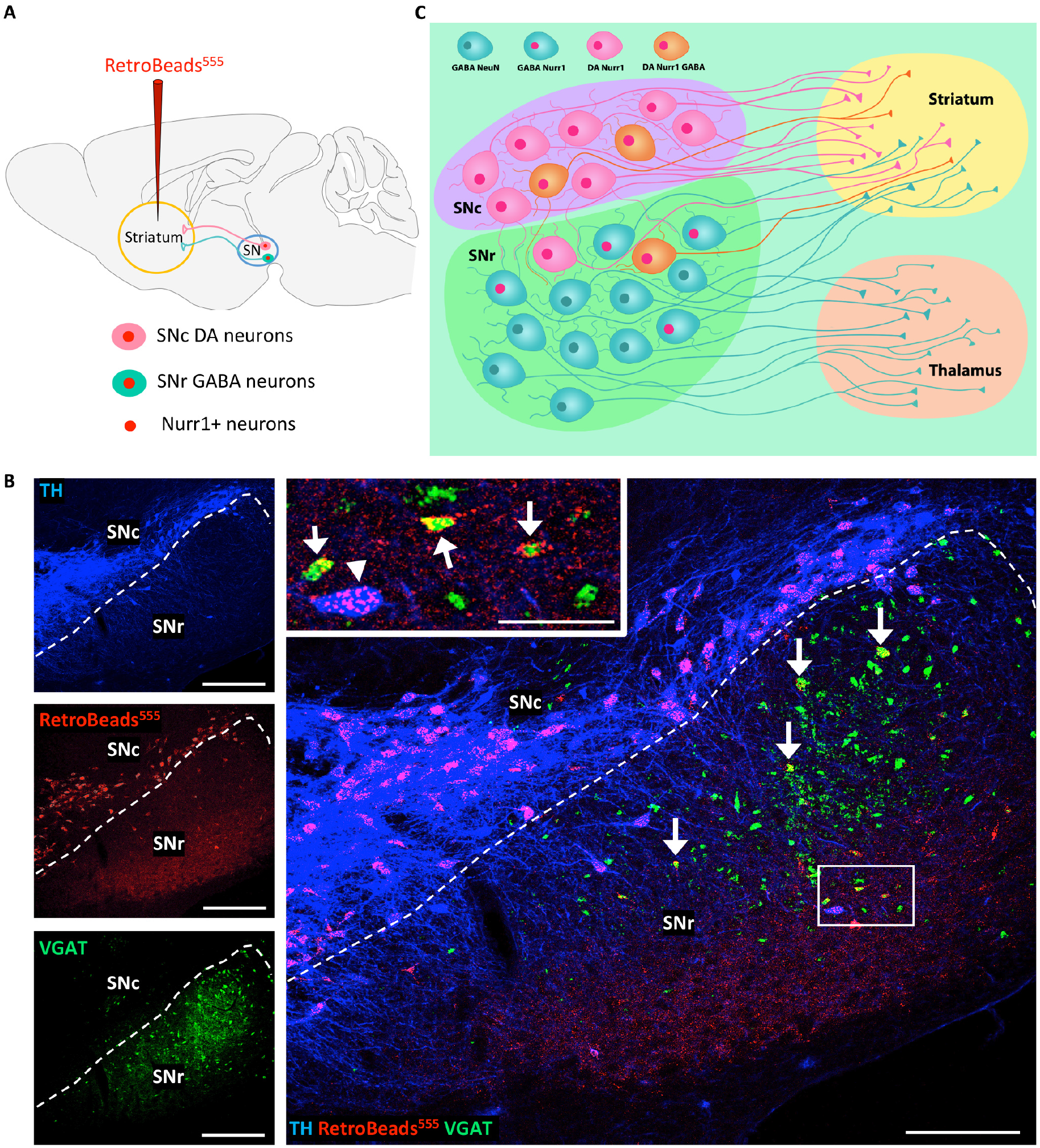
Retrograde tracing revealed the pool of SNr GABAergic neurons projecting to the striatum. **A**, Schematic diagram illustrating retrograde tracing with RetroBeads (555 nm) injected in the striatum and transported from the neuronal terminals in the striatum back to their somata in the SN. **B**, Confocal images of representative coronal sections through the SN of adult (P60) VGAT-ZsGreen mice. RetroBeads were detected in the somata of both TH+ (inset, arrowhead) and VGAT+ (arrows) neurons in the SNr, revealing the connectivity of GABAergic SNr-to-striatum projection neurons. Scale bars = 200 μm, 50 μm (inset). **C**, Schematic diagram of neuronal projections from SN to striatum and thalamus. The SNc is densely packed with DAergic nigrostriatal neurons, a fraction of which co-expresses GABA. The SNr includes GABAergic cells, a fraction of which expresses Nurr1 (pink nuclei) and projects to striatum, sharing the same target with SNc DAergic neurons.

Our IHC data showed that DAergic neurons in the SNc display a rich dendritic arborization extending into the SNr (Fig. S3, arrows). Such anatomical connectivity is in agreement with previous studies demonstrating that SNr GABAergic neurons can be electrically excited by direct activation of D_1_ and D_5_ receptors mediated by DA release from SNc DAergic dendrites^49^ and nicotine-mediated activation of nAChRs^25,26^.

### Nurr1 upregulation is sufficient to ameliorate PD-related locomotor deficits and decrease the number of hα-syn+ neurons in the SN

Because chronic nicotine exposure attenuated PD-related locomotor deficits and induced an increase in the expression of Nurr1 in the SNr (Fig. 1), we next investigated the effects of expanding the pool of Nurr1-expressing neurons in the SN on locomotor performance in PD mice. We injected a pan-neuronal viral vector (AAV.TRMS.Nurr1) into the SN of transgenic PD mice to upregulate Nurr1 when robust accumulation of hα-syn in TH+ neurons was already present (P111). Successful hα-syn expression was verified by IHC. This experiment included three groups: hα-syn-negative mice without viral injection (hα- syn-), hα-syn+ mice injected with a viral vector expressing GFP (hα-syn+_AAV.GFP), and hα-syn+ mice injected with AAV.TRMS.Nurr1 (hα-syn+_AAV.TRMS.Nurr1). Animals were given 8 weeks to recover from surgery and reach peak transgene expression^50^ before being tested for locomotor behavior at P167. Only animals displaying transgene expression in the SN, which was assessed later by IHC (Fig. 6C, F), were included in the BPM analysis. In agreement with our previous findings (Fig.1), hα-syn+ mice injected with AAV.GFP showed a number of motor deficits when compared to hα-syn- mice (Fig. 6A-B). Mixed model ANOVA showed significant differences (Fig. 6A) for distance traveled (time x group interaction: F_(6,85)_=2.258, p<0.05, group main effect: F_(2,30)_=3.848, p<0.05), transitions (group main effect: F_(2,29)_=5.209, p<0.05), and entries to center (time x group interaction: F_(6,88)_=4.176, p<0.001). Strikingly, hα-syn+ mice overexpressing Nurr1 did not show locomotor deficits, exhibiting a level of behavioral performance similar to hα-syn- mice. In the latter half (20-40 min) of the testing session, locomotor differences (Fig. 6B, one-way ANOVA) were observed in distance traveled (F_(2,30)_=5.445, p<0.01), transitions (F_(2,29)_=6.833, p<0.01), and entries to center (F_(2,28)_=9.293, p<0.001). Behavioral improvements were significant in Nurr1-treated animals as compared to the GFP control (Fig. 6B, all three measures, Bonferroni’s Multiple Comparisons: hα-syn+_AAV.GFP vs hα-syn+_AAV.Nurr1, p<0.05).

**Figure 6.**
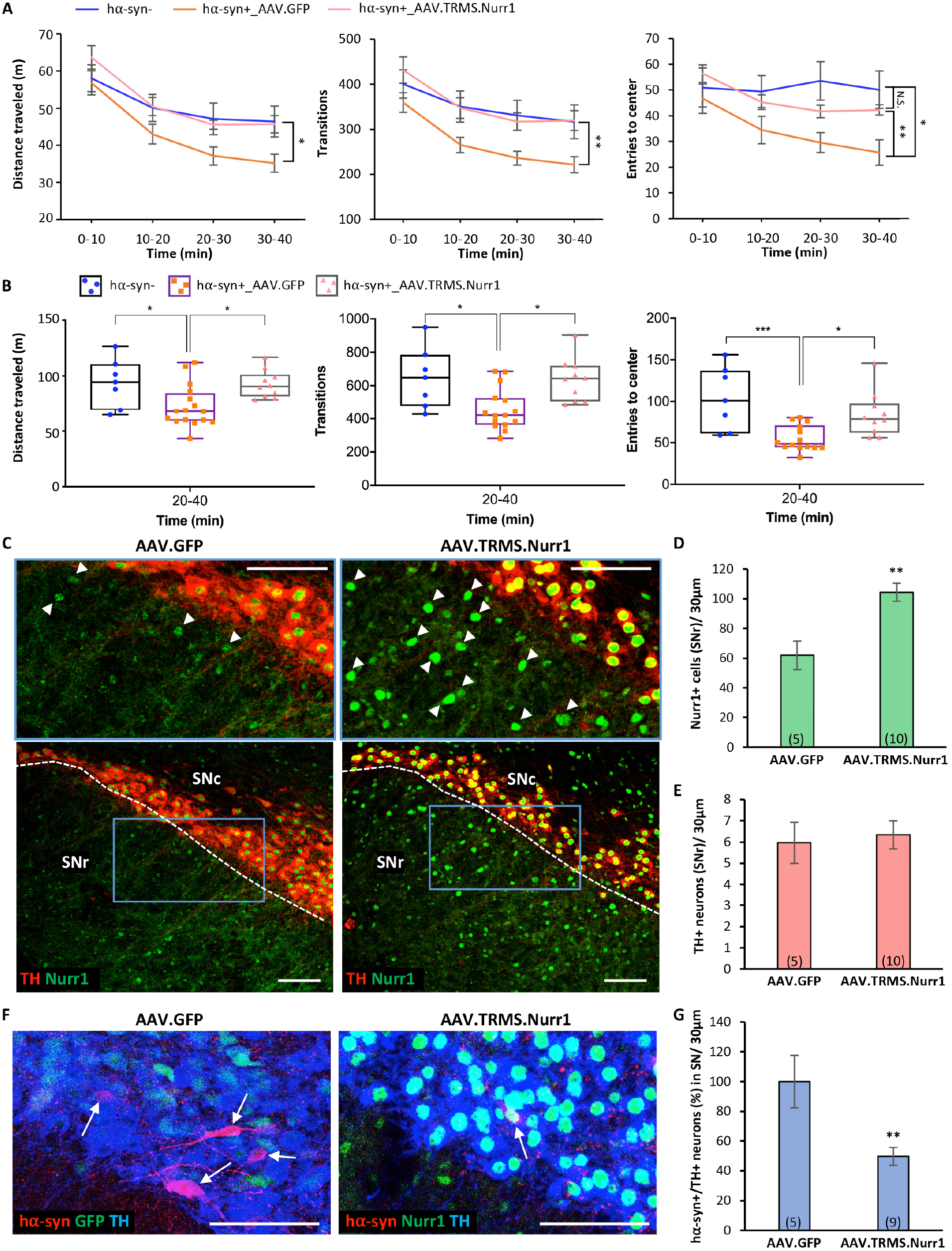
Nurr1 overexpression is sufficient to ameliorate PD-related locomotor deficits and decreases the number of hα-syn+ neurons in the SNc. **A,** Locomotor activity (BPM) measured from 0 to 40 min of the testing session showed that pan-neuronal Nurr1-overexpression ameliorated PD-related locomotor deficits exhibited by hα-syn+_AAV.GFP mice (mixed model ANOVA for distance traveled: time x group interaction, F_(6,85)_ = 2.258, p<0.05, main effect of group, F_(2,30)_ = 3.848, p<0.05, Bonferroni’s Multiple Comparisons: hα-syn+_AAV.GFP vs hα-syn+_AAV.Nurr1 p<0.05 at 20-30 min and 30-40 min; transitions: main effect of group: F_(2,29)_ = 5.209, p<0.05, Bonferroni’s Multiple Comparisons: hα-syn+_AAV.GFP vs hα-syn+_AAV.Nurr1 p<0.05 at 10-20, 20-30 min, and p<0.01 at 30-40 min; entries to center: time x group interaction, F_(6,88)_ = 4.176, p<0.001, Bonferroni’s Multiple Comparisons: hα-syn+_AAV.GFP vs hα- syn- p<0.05, vs hα-syn+_AAV.Nurr1 p<0.01 at 30-40 min. Every measure shows a main effect of time, p<0.0001. Graphs show mean ± SEM: *p<0.05, **p<0.01, N.S., not significant. The number of animals for each group is: hα-syn- (N=7), hα-syn+_AAV.GFP (N=17), hα-syn+_AAV.Nurr1 (N=10). **B**, Locomotor measures (distance traveled, transitions, and entries to center) plotted for 20-to-40 min interval of the BPM testing session shown in **A** revealed a significant AAV.Nurr1-mediated rescue of the behavioral deficits displayed by hα-syn+_AAV.GFP mice. One-way ANOVA for distance traveled: F_(2,30)_ = 5.445, p<0.01, Bonferroni’s Multiple Comparisons: hα-syn+_AAV.GFP vs hα-syn- p<0.05, vs hα-syn+_AAV.Nurr1 p<0.05; transitions: F_(2,29)_ = 6.833, p<0.01, Bonferroni’s Multiple Comparisons: hα- syn+_AAV.GFP vs hα-syn- p<0.05, vs hα-syn+_AAV.Nurr1 p<0.05; entries to center: F_(2,28)_ = 9.293, p<0.001, Bonferroni’s Multiple Comparisons: hα-syn+_AAV.GFP vs hα- syn- p<0.001, vs hα-syn+_AAV.Nurr1 p<0.05). Graphs show all data points with medians and interquartile range. *p<0.05, **p<0.01, ***p<0.001. **C**, Confocal images of representative SN sections showing enhanced Nurr1 immunoreactivity (arrowheads) in mice injected with pan-neuronal Nurr1 viral vector (AAV5.TRMS.Nurr1) compared to mice injected with AAV.GFP (AAV5.TRMS.GFP). Scale bars = 100 μm. **D-E**, Quantification of IHC preparations in **C** showed that AAV.TRMS.Nurr1 injection increased the number of Nurr1+ cells in the SNr (**D**, t_(13)_=3.86, p<0.01) but was not sufficient to lead to an increase in TH expression in the SNr (**E**). Graphs show mean ± SEM: **p<0.01. The number of animals is annotated in parenthesis for each condition. **F**, Confocal images of representative SN sections showing hα-syn+/TH+ neurons (arrows) in AAV.GFP- and AAV.TRMS.Nurr1-injected mice. Scale bars = 50 μm. **G**, Quantification (%) of IHC preparations shown in **F** indicates that AAV.TRMS.Nurr1 injection decreased the number of hα-syn+/TH+ neurons in the SN (t_(12)_=3.33, p<0.01). Graphs show mean ± SEM: **p<0.01. The number of animals is annotated in parenthesis for each condition.

After behavioral testing, brain tissue was processed by IHC with hα-syn, TH, and Nurr1 markers to investigate the effects of Nurr1 overexpression on these molecular phenotypes in the SN. We confirmed that mice injected with AAV.TRMS.Nurr1 displayed enhanced Nurr1 immunoreactivity (Fig. 6C, arrowheads), exemplified as an increased number of Nurr1+ cells in SNr (Fig. 6D, mean±SEM: AAV.GFP = 62±10, AAV.TRMS.Nurr1 = 104±6, t_(13)_=3.86, p<0.01). While the number of TH+ neurons remained unchanged (Fig. 6E, mean±SEM: AAV.GFP = 6.0±1.0, AAV.TRMS.Nurr1 = 6.3±0.7), the number of hα-syn+/TH+ neurons in the SN (Fig. 6F, arrows) of hα-syn+ mice injected with AAV.TRMS.Nurr1 was 50% lower than AAV.GFP-injected ones (Fig. 6G, mean±SEM: AAV.GFP = 100±18 %, AAV.TRMS.Nurr1 = 50±6 %, t_(12)_=3.33, p<0.01), suggesting that Nurr1 overexpression resulted in a neuroprotective effect against hα-syn toxicity.

### Selective Nurr1 upregulation in GABAergic cells is not sufficient to induce a TH phenotype

Because the fraction of SNr GABAergic neurons projecting to the striatum represents a reserve pool that can acquire Nurr1 and TH phenotypes in response to nicotine-mediated activation, we tested whether Nurr1 upregulation alone, exclusively targeted to SN GABAergic cells, could induce TH plasticity. We unilaterally injected a Cre-dependent Nurr1 viral vector (AAV.FLEX.Nurr1) into the SN of VGAT-Cre mice (P60). The contralateral uninjected side was used as control. Nurr1 immunoreactivity in the AAV.FLEX.Nurr1-injected side, as compared to control (Fig. 7A, arrowheads), showed robust vector transduction. A quantitative analysis showed that the increase in the number of Nurr1-expressing VGAT+ cells (Fig. 7B, mean±SEM: control = 54±3, AAV.FLEX.Nurr1 = 98±12, t_(7)_=2.96, p<0.05) was not paralleled by an increase in the number of TH+ neurons in the SNr (Fig. 7C, mean±SEM: control = 11±1, AAV.FLEX.Nurr1 = 12±1), indicating that Nurr1 upregulation alone was not sufficient to induce the acquisition of TH phenotype by SNr GABAergic neurons.

**Figure 7.**
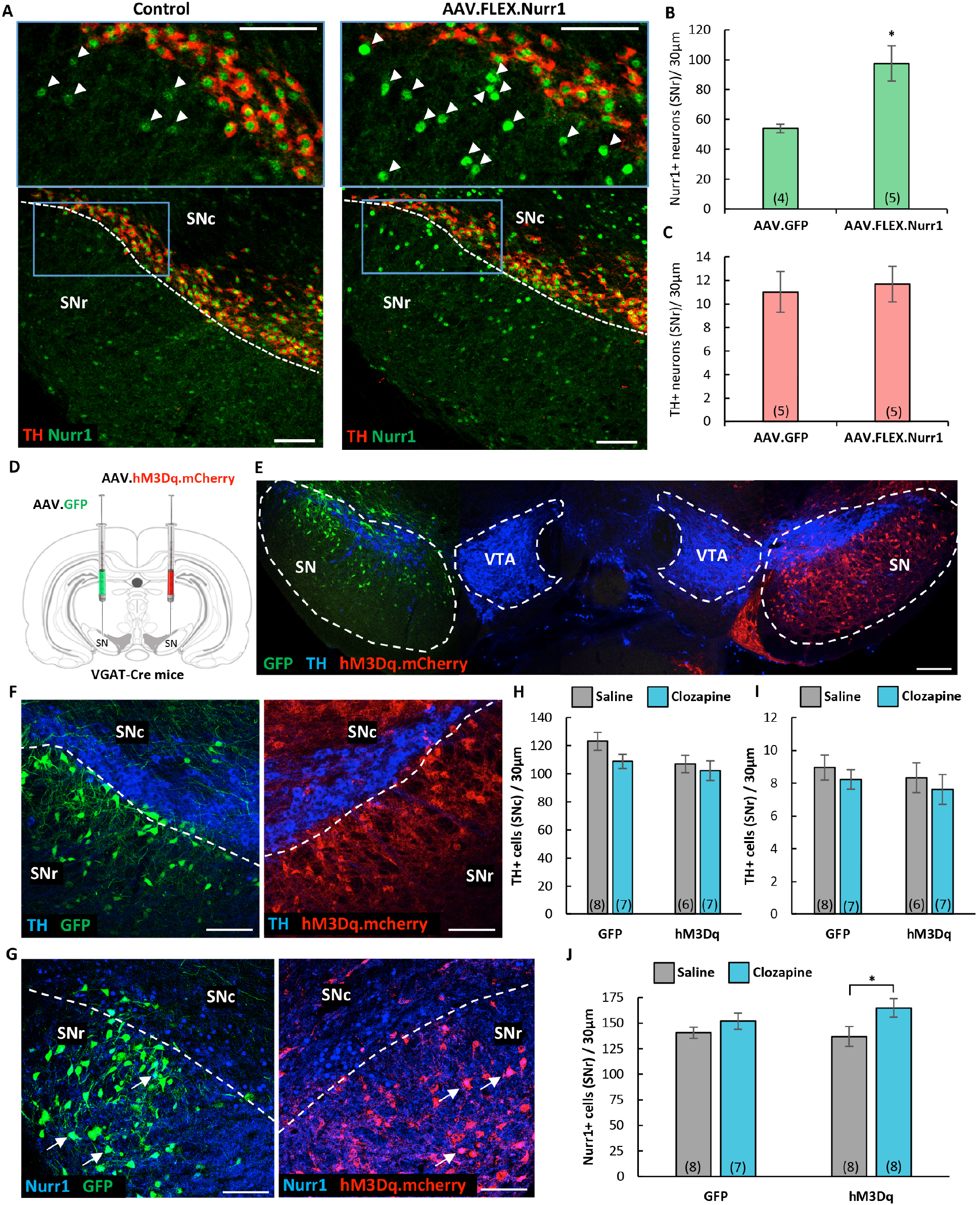
Selective Nurr1 overexpression does not elicit a TH phenotype while chronic activation of GABAergic cells is sufficient to induce Nurr1. **A**, Confocal images of representative SN sections showing enhanced Nurr1 immunoreactivity (arrowheads) on the side of the brain injected with Cre-dependent AAV Nurr1 miRNA (AAV.FLEX.Nurr1), when compared to the contralateral uninjected side (control). Scale bars = 200 μm. **B-C**, Quantification of IHC preparations shown in A indicates that AAV.FLEX.Nurr1 injection increased the number of Nurr1+ cells in the SNr (**B**, t_(7)_=2.96, p<0.05) but was not sufficient to lead to an increase in TH expression in the SNr (**C**). Graphs show mean ± SEM: *p<0.05. The number of animals is annotated in parenthesis for each condition. **D**, Schematic representation of the viral strategy adopted to chemogenetically activate VGAT+ neurons of the SN of VGAT-Cre mice by unilaterally injecting a Cre-dependent DREADD (hM3Dq).mCherry AAV and transfecting the contralateral side with a Cre-dependent AAV.GFP reporter (control). **E**, Composite image of a representative section (30 μm) through the SN and VTA showing selective expression of Cre-dependent mCherry-tagged excitatory DREADD (hM3Dq) virus within the SN GABAergic neurons on the right side and Cre-dependent GFP virus transfected on the left side of a VGAT-Cre mouse brain, along with TH immunofluorescent staining. Scale bars = 200 μm. **F-G**, Confocal images of representative SN sections showing TH (**F**) and Nurr1 (**G**, arrows) immunoreactivity along with GFP and hM3Dq.mCherry labeling. Scale bars = 100 μm. **H-I**, Quantification of TH+ neurons of IHC preparations shown in **F**,**G** (**H**, SNc, two-way ANOVA, main effect of hM3Dq, F_(1,43)_ = 4.49, p<0.05; **I**, SNr, unaffected by chemogenetic activation of VGAT+ neurons). Graphs show mean ± SEM. The number of animals is annotated in parenthesis for each condition. **J**, Quantification of IHC preparations shown in **G** revealed that DREADD (hM3Dq)-mediated activation of GABAergic cells induced an increase in Nurr1 expression (two-way ANOVA, main effect of clozapine, F_(1,14)_ = 5.325, p<0.05, Bonferroni’s Multiple Comparisons: hM3Dq/saline vs hM3Dq/clozapine, p<0.05). Graph shows mean ± SEM: *p<0.05. The number of animals is annotated in parenthesis for each condition.

### Chemogenetic activation of SN GABAergic neurons is sufficient to induce the acquisition of Nurr1 but not TH phenotype

Given that chronic alteration of neuronal activity^44,51,52^ and nicotine-exposure^43,53^ can elicit TH plasticity within non-DAergic interneurons of the activated network, we utilized a chemogenetic approach with DREADDs (designer-receptors-exclusively-activated-by-designer-drugs), to test whether chronic depolarization of GABAergic neurons was sufficient to induce TH plasticity. To this end, we unilaterally injected a viral vector carrying a Cre-dependent excitatory mCherry-DREADD (hM3Dq) construct into the SN of adult VGAT-Cre mice (P60) and injected the contralateral side with a Cre-dependent AAV vector expressing GFP as a control (Fig. 7D). One month after viral infusion (P90), mice clearly exhibited GFP and hM3Dq.mCherry expression ipsilateral to the side of infusion in the SN (Fig. 7E). At this point, we began our DREADD-activation protocol by administering 0.01 mg/kg clozapine or saline as control (i.p., twice daily) for 14 days. Quantification of TH immunoreactivity (Fig. 7F) showed that chronic activation of GABAergic cells was not sufficient to induce an increase in the number of TH-expressing neurons in either SNc or SNr (Fig. 7H-I). However, DREADD-mediated activation was sufficient to elicit a significant increase in the number of Nurr1+ neurons exclusively in the DREADD-injected hemisphere (Fig. 7G, arrows; Fig. 7J, mean±SEM: GFP/saline = 141±5, GFP/clozapine = 152±8, hM3Dq/saline = 137±10, hM3Dq/clozapine = 165±9, two-way ANOVA, clozapine main effect: F_(1,14)_ = 5.325 p<0.05, Bonferroni’s Multiple Comparisons: hM3Dq/saline vs hM3Dq/clozapine, p<0.05).

## DISCUSSION

Our findings show that chronic nicotine exposure attenuates locomotor deficits in a human-α-syn-expressing mouse model of PD^41^ and primes a GABAergic neuronal pool in the SNr to a novel form of neuroplasticity culminating in the acquisition of the TH phenotype. Nicotine activation of nAChRs in the nigrostriatal pathway elicits an increase in calcium influx^22–24^ and induces neuronal depolarization^20,21^. Previous studies have reported on the dense distribution of nAChRs in both DAergic and GABAergic neurons in the SN^25,26^, suggesting a potential activity-dependent mechanism in the regulation of DAergic circuits in nicotine-mediated protection against PD^18,54–56^. Specifically, 99% of SNr GABAergic neurons express both α_4_* nAChR readily available for nicotine activation^25,26^ as well as DA D_1_ and D_5_ receptors which are tonically excited by dendritically released DA from the SNc DAergic neurons, forming a relatively short SNc- to-SNr DAergic pathway^49^. The dendro-dendritic connectivity and the specific receptor expression displayed by descending SNc DAergic dendrites into the SNr GABAergic neuropil (Fig. S3) provide the opportunity for the nigrostriatal circuit to signal common instructions to both the DAergic and the GABAergic pathways when DA function needs a physiological boost. These conditions have been shown to be a requirement for activity-dependent recruitment of non-DAergic neurons to a DAergic phenotype^44^. Chronic nicotine exposure could elicit the recruitment of SNr GABAergic neurons to Nurr1 and TH phenotypes through at least two potential activity-dependent signaling mechanisms: (a) nicotine directly activates the α_4_* nAChRs localized on the SNr GABAergic neurons; (b) as nicotine activates SNc DAergic neurons via nAChRs, the DA released from the dendrites activates SNr GABAergic neurons through D_1_ and D_5_ receptors. Both mechanisms could, in principle, initiate the calcium-mediated reprogramming required to induce the TH phenotype in the SNr GABAergic neurons, as previously found in neurons of the SNc^57^.

Electrical activity and calcium signaling have significant roles in regulating various forms of neuroplasticity, including priming neurons with the molecular memory of early drug exposure^53^ and neurotransmitter reprogramming^52,58,59^. Sustained alteration in circuit activation by either experimental manipulation or natural sensory stimuli can induce neurotransmitter plasticity in the mature brain, affecting behavior^51,53, 60–62^. While SNr GABAergic neurons undergo a significant upregulation of α_4_* nAChR subtype in response to chronic nicotine exposure, the level of these receptors in SNc DAergic neurons remains unchanged^26^. Selective upregulation of α4* nAChR level in the SNr might bring the level of calcium transients in these neurons to a threshold sufficient to signal and initiate neurotransmitter plasticity in response to chronic nicotine exposure, providing another layer of specificity in the recruitment of GABAergic neurons of the SNr and not the SNc to nicotine-mediated TH plasticity.

We identified a reserve pool of Nurr1-expressing GABAergic neurons in the SNr that undergoes nicotine-mediated TH respecification; a phenomenon that might represent a layer of functional protection against PD. Our findings provide an important parallel to previously reported phenotypic shift of pre-existing GABAergic neurons to express TH in adult macaques following treatment with MPTP, a neurotoxin that induces DA depletion mimicking PD^63^. Importantly, we confirmed by retrograde tracing that part of the nigrostriatal projection originates from SNr GABAergic neurons and demonstrated that these neurons share the same target as DAergic neurons in the SNc. Therefore, these SNr GABAergic neurons could serve the role as a reserve pool that could gain the DA-synthesizing enzyme and potentially rescue the DAergic loss of function caused by neurodegeneration of SNc DAergic neurons. Given that this form of TH plasticity also occurs in the SN of primates^64^ in response to DAergic neuron loss, understanding the mechanism of nicotine-induced TH respecification in the SNr has tremendous translational value in the constant search for new approaches aimed at replenishing DA function in PD.

As chronic nicotine exposure leads to improved locomotion in hα-syn+ mice and concomitantly increases the number of Nurr1+ cells in the SNr, we further investigated the effect of the induced upregulation of Nurr1 expression in the SN of hα-syn+ mice. We found that Nurr1 overexpression was sufficient to ameliorate PD-related locomotor deficiencies. This is in agreement with previous studies highlighting the role of Nurr1 in pathogenesis of PD and its potential as a therapeutic target^36,37^. Our results here show, for the first time, that Nurr1 overexpression elicits protection against PD-related locomotor dysfunctions through a reduction of the number of hα-syn-expressing TH+ neurons. Given the established neurodegenerative effects of abnormal α-synuclein^65^ and the therapeutic effect of Nurr1^36,37^, chronic nicotine exposure might slow down the etiology of neurodegeneration by reducing and attenuating α-syn toxicity.

Our chemogenetic approach implemented to selectively and chronically depolarize GABAergic neurons was not sufficient to induce TH plasticity in these neurons; however, it revealed a way to experimentally induce an expansion of the reserve pool of Nurr1+ neurons in the SNr that is available for recruitment to TH phenotype acquisition. Neither selective Nurr1 upregulation nor chronic activation of GABAergic neurons alone recapitulated TH respecification observed in the SNr of mice chronically exposed to nicotine. The multi-factorial nature of the nicotine stimulus includes simultaneous upregulation and chronic activation of nAChRs which in turn could determine the exact calcium dynamics regulating a cassette of DAergic genes encoding Nurr1 and TH. The α_4_* nAChR subtypes, expressed in most SNr GABAergic neurons^26^, are among the most nicotine-sensitive subtypes^66^ and are upregulated by chronic nicotine^67^. The upregulated nicotinic receptors retain functions leading to increased high-affinity binding sites for nicotine and potentiation of the nicotinic response in brain synaptosomes^68–70^. As chronic nicotine upregulates nAChRs, more SNr GABAergic neurons might become activated via nAChR-mediated calcium signaling. Calcium-mediated activity regulates Nurr1 expression^32^ and membrane depolarization promoting DAergic differentiation by increasing TH+ neurons and *TH* mRNA in non-DAergic cells^51^. The calcium signaling threshold needed for recruiting the DAergic phenotype^53,71^ or other homeostatic forms of NT plasticity^52^ varies by cell types. This might explain why TH respecification elicited by chronic nicotine exposure was not recapitulated by chemogenetic activation.

Future studies will uncover all key players required for a combinatorial manipulation that would successfully induce SNr GABAergic neurons to acquire the TH phenotype. Since the SNr GABAergic fraction of the nigrostriatal pathway is completely spared by PD-associated neurodegeneration, the gain of the DAergic phenotype could in principle replenish DA in the striatum. Establishing effective manipulations targeted to induce TH plasticity in GABAergic neurons of the nigrostriatal pathway could represent a paradigm shift in developing a novel approach for PD treatment.

## ACKNOWLEDGEMENTS

We thank Dr. H. Cai for generously providing the initial breeding pair of the *Pitx3-IRES2-tTA* mice, and Dr. M. Ulivieri for critical comments on the manuscript.

## Author Contributions

D.D. planned the project; I.L. designed and carried out the experiments and performed data analysis; B.R. performed viral injections; M.K. assisted with confocal image acquisition and cell quantification; S.P. contributed to generate and analyze the BPM data; F.P.M. provided the Nurr1 viral vectors; I.L. and D.D. wrote the manuscript.

