## Supplement for "Nicotine-mediated recruitment of GABAergic neurons to a dopaminergic phenotype attenuates motor deficits in alpha-synuclein Parkinson’s model"

### METHODS AND MATERIALS

#### Animals

Mice were housed in groups of two to five in a standard 12h dark: 12h light cycle and fed with regular diet *ad libitum*. Male and female transgenic mice used in this study were genotyped by polymerase chain reaction (PCR) analysis of ear biopsies. All experiments were conducted in accordance with protocols approved by the UCSD Institutional Animal Care and Use Committee (IACUC) and guidelines of the American Association for the Accreditation of Laboratory Animal Care and National Research Council's Guide for the Care and Use of Laboratory Animals.

<u>GAD67-GFP knock-in mice</u>

The GAD67-GFP knock-in mouse line was kindly provided by Dr. Yuchio Yanagawa, Gunma University Graduate School of Medicine, Japan. Mice were heterozygous for insertion of the gene encoding green fluorescent protein (GFP) under the control of the GAD67 gene promoter[1]. They were used to label the inhibitory GABAergic neurons in the

SN by enhancing the GFP signal with an anti-GFP antibody. Adult mice (P60) weighing 25-35g were used in this study.

VGAT-Cre knock-in mice

VGAT-IRES-Cre knock-in mice (STOCK *Slc32a1*$^{tm2(cre)Lowl}$/, Jackson stock 016962), expressing Cre under the control of VGAT regulatory elements, were bred in our vivarium and used for chemogenetics. To induce the expression of the ZsGreen label in GABAergic cell bodies, VGAT-Cre mice were bred with reporter mice (B6.Cg-*Gt(ROSA)26Sor*$^{tm6(CAG-ZsGreen1)Hze}$/J, Jackson stock 007906) that express CAG promoter-driven enhanced green fluorescent protein (ZsGreen1) following Cre-mediated recombination.

*Pitx3-IRES2-tTA/tetO-A53T* double transgenic mice

The *Pitx3-IRES2-tTA/tetO-A53T* double transgenic mouse line, which expresses mutant (*SNCA*A53T*) human α-synuclein in midbrain dopaminergic neurons, was previously characterized[2] and generously provided by Dr. Cai at NIH. By crossing the driver line, *Pitx3-IRES-tTA* mice (B6.129(FVB)-*Pitx3*$^{tm1.1Cai}$/J, Jackson stock 021962), with the responder line, *tetO-A53T*, which encodes a human α-synuclein mutant gene under the control of a *tetO* promoter (STOCK Tg(tetO-SNCA*A53T)E2Cai/J, Jackson stock 012442), the expression of A53T α-synuclein in the SN dopaminergic neurons was driven using a binary tetracycline-dependent “tet-off” inducible gene expression system. Breeders were given doxycycline (DOX)-containing (200mg/kg) food pellets (Bio-Serv), in place of a regular diet, to suppress transgene expression from early embryonic stages through weaning (P21). Adult mice, weighing 20-30g, were used to investigate either the accumulation of human α-synuclein (P30 through P180) or the effects of nicotine exposure. Mice of the responder line *tetO-A53T* were used as controls (hα-syn−).

**Stereotactic injections**

Mice were anesthetized with 3% isoflurane and placed in a stereotactic apparatus (David Kopf Instruments, Model 900HD Motorized Small Animal Stereotaxic). Brain injections were performed during a continuous flow of 1% isoflurane.

<u>Chronic chemogenetic activation via designer receptor exclusively activated by designer drugs (DREADD)</u>

Adult VGAT-Cre mice (P60 or P90) underwent bilateral stereotactic injections (500nL/side, $1.0 \times 10^{13}$ GC/mL) into the SN (AP = -3.08 mm, L = ±1.38 mm, DV = -4.66 mm) with a control virus (AAVDJ.Syn1.DIO.eGFP, Salk Institute, La Jolla, CA) on one side and the excitatory DREADD receptor-encoding virus (AAV5.hSyn.DIO.hM3Dq.mCherry, Addgene, Watertown, MA) on the contralateral side.

<u>Nurr1 Overexpression</u>

Genomes for overexpression were the same as used previously[3] where Nurr1 or GFP expression is driven by the CAG promoter. Cre-dependent vectors contained the same cassette, however, the transgene is floxed and inverted in the un-induced state. AAVs were produced as described previously[4]. Briefly, 293T cells were co-transfected with genome-encoding plasmids as well as plasmids carrying cap genes (AAV5 or AAV9) or helper functions. Following harvest, vectors were purified using an iodixanol gradient followed by dialysis and concentration in modified PBS. Titers were determined using ddPCR.

For pan-neuronal Nurr1 overexpression in the SN (AP = -3.08 mm, L = ±1.38 mm, DV = -4.66 mm), adult (P111) hα-syn+ mice were injected with either AAV5.TRMS.Nurr1[3] (300 nL, $1.0 \times 10^{13}$ GC/mL) or with AAV.TRMS.GFP (300 nL, $1.6 \times 10^{13}$ GC/mL) control virus. For Cre-dependent Nurr1 overexpression in GABAergic neurons of the SN, adult (P90) VGAT-Cre mice were injected with AAV9.FLEX.Nurr1 (300 nL, $1.0 \times 10^{13}$ GC/mL); the control group was injected with AAV9.FLEX.GFP (300 nL, $1.0 \times 10^{13}$ GC/mL).

To allow diffusion of the injected virus, the injection needle remained in place for 8 minutes before removal. After surgery, mice were injected with 0.1mg/kg/24h buprenorphine as analgesia. Viral incubation occurred for 4-6 weeks after surgery. Viral vectors were generously provided by Dr. Fredric Manfredsson, Barrow Neurological Institute, AZ[5].

Retrograde tracing

Fluorescent RetroBeads (80 nL, 555nm, LumaFluor, Inc., Durham, NC) was unilaterally injected in the striatum (AP = −0.20 mm, L = ±2.60 mm, DV = −3.00 mm) of VGAT-ZsGreen mice. Mice were sacrificed after 10 days of recovery to allow adequate time for retrograde transport of RetroBeads from the striatal terminals to the soma of SN neurons.

**Drug treatment**

Chronic nicotine exposure

Two groups of adult GAD67-GFP (P60), VGAT-Cre (P60), or hα-syn+ (P111) mice underwent chronic nicotine exposure for two weeks. Drinking water was replaced with a solution of 50mg/L nicotine in 1% saccharin (nicotine-treated group) or with 1% saccharin solution (control condition). Animals were sacrificed after the two-week treatment. The amount of fluid intake was measured daily throughout the experiment (Fig. S2A-B); the initial and final body weight of the mice were also measured (Fig. S2C).

Chronic chemogenetic activation via DREADD

Clozapine (0.01 mg/kg; MP Biomedicals, Santa Ana, CA) was dissolved in 0.1% dimethyl sulfoxide (DMSO) in sterile saline or vehicle (sterile saline) was administered intraperitoneally (IP) to VGAT-Cre mice twice daily for 14 days.

**Immunohistochemistry and *in situ* hybridization**

For immunohistochemistry (IHC), mice were anesthetized with a ketamine/xylazine cocktail (10mg/kg) delivered via i.p. injection. Animals were transcardially perfused with room-temperature phosphate buffered saline (1X PBS) followed by ice-cold 4% paraformaldehyde (PFA). Brains were incubated 4% PFA overnight at 4°C and transferred to 30% sucrose for 48-72 hours until sunk. Brains were then serially sectioned at 30 µm using a Leica microtome (SM 2010R) and collected in PBS.

For RNAscope *in situ* hybridization (ISH), mice were anesthetized with ketamine/xylazine (10mg/kg) via i.p. injection and euthanized by decapitation. Brains were removed, snap-frozen with powdered dry ice, and stored in −80°C. Brains were serially cut at 20 µm using a Leica cryostat (CM 1860) and mounted directly onto glass slides. Slides were stored at −80°C before RNAscope processing.

All washes and incubations were performed with gentle rotation. Antibodies (listed on Tabel 1) were diluted in blocking solution (1X PBS containing 5% normal horse serum and 0.3% Triton X-100).

For immunofluorescence, free-floating brain sections were washed 3 times for 10 minutes (3 x 10') in PBS, incubated in blocking solution for 30' to 1 hour and with primary antibodies in blocking solution overnight at 4°C. The following day, sections were washed 3 x 10' in PBS, then incubated in secondary antibodies in blocking solution for 1 hour at room temperature, washed 3 x 10' in PBS, mounted on a positively charged glass slide (Fisherbrand Superfrost Plus) with 0.2% gelatin in PBS, coverslipped with mounting medium (Fluoromount-G®, SouthernBiotech) with or without DRAQ5 (1µm/mL, BioStatus), and sealed with nail polish for permanent storage and imaging.

For colorimetric DAB (3,3-Diaminobenzidine)-based IHC, free-floating sections were washed three times (10 minutes per wash) in PBS, then incubated in ABC (Avidin-Biotin Complex) solution (1X PBS containing 0.3% Triton X-100, 2% NaCl, and 1% of Reagents A and B from the VECTASTAIN® ABC kit) for one hour, washed again (3 x 10 minutes), and incubated in fresh DAB solution (25mg/mL) for approximately three minutes depending on the speed of the reaction. Sections were rinsed twice quickly and washed for 20 minutes in PBS before mounting on glass slides. Sections on slides were dried in a fume hood, then defatted in 1:1 chloroform:ethanol solution for two hours, progressively rehydrated in 100% ethanol, 95% ethanol, and distilled water. Sections were then counterstained in 0.1% cresyl violet solution for 30 minutes, rinsed quickly in distilled water, dehydrated in 95% ethanol for three minutes, in 100% ethanol twice for five minutes each, cleared (2 x 5 min) in Xylenes (brand). Slides were coverslipped with permanent mounting medium Cytoseal™ 60 (Thermo Scientific). Images were acquired using Hamamatsu Nanozoomer 2.0HT Slide Scanner. The quantification was performed by unbiased stereology (using a Leica DM4 B microscope and Stereologer2000 software). RNAscope in situ hybridization to detect and label TH and Nurr1 mRNA transcipts was performed following manufacturer instructions (Advanced Cell Diagnostics). Sections were counterstained with DAPI and slides coverslipped using Fluoromount-G mounting medium. Images were acquired at 20X magnification with a confocal microscope (Leica TCS SPE).

| **Primary Antibody** | **Manufacturer, Catalog #** | **Dilution** |
|---|---|---|
| Chicken anti GFP | Invitrogen, A10262 | 1:500 |
| Chicken anti GFAP | Invitrogen, AB5541 | 1:1000 |
| Goat anti Foxa2 | Boster, A01032 | 1:250 |
| Guinea Pig anti NeuN | Millipore Sigma, ABN90 | 1:1500 |
| Mouse anti TH | Millipore Sigma, MAB318 | 1:1000 |
| Mouse anti hα-syn | Santa Cruz, sc-12767 | 1:500 |
| Rabbit anti Nurr1 | Santa Cruz, sc-990 | 1:300 |
| Sheep anti TH | Novus, NB300-110 | 1:1000 |
| **Secondary Antibody** | **Manufacturer, Catalog #** | **Dilution** |
| Donkey anti chicken FITC | Invitrogen, SA1-72000 | 1:300 |
| Donkey anti chicken Alexa Fluor 647 | Millipore Sigma, AP194SA6 | 1:500 |
| Donkey anti goat Alexa Fluor 488 | Invitrogen, A11055 | 1:500 |
| Goat anti guinea pig Alexa Fluor 488 | Abcam, ab150185 | 1:500 |

| | | |
|---|---|---|
| Goat anti guinea pig Alexa Fluor 594 | Invitrogen, A11076 | 1:500 |
| Donkey anti mouse Alexa Fluor 488 | Invitrogen, A21202 | 1:500 |
| Donkey anti mouse Alexa Fluor 555 | Invitrogen, A31570 | 1:500 |
| Donkey anti mouse Alexa Fluor 647 | Invitrogen, A31571 | 1:500 |
| Donkey anti rabbit Alexa Fluor 488 | Invitrogen, A21206 | 1:500 |
| Donkey anti rabbit Alexa Fluor 555 | Invitrogen, A31572 | 1:500 |
| Goat anti rabbit Alexa Fluor 647 | Invitrogen, A21244 | 1:500 |
| Donkey anti sheep Alexa Fluor 647 | Invitrogen, A21448 | 1:500 |

**Table S1 – List of Antibody and dilution ratio.**

**Quantification of immunoreactivity**

For immunofluorescence, sections including the rostrocaudal extent of the substantia nigra were collected and stained for TH, Nurr1, Foxa2, NeuN, and GFP. Images were acquired using a Leica TCS SPE confocal microscope and used for quantification of neurons with various IHC markers. Maximized fluorescence final images were obtained from a total of 11 Z-stacked layers 2 μm away from each other. Cells were counted by an investigator blind to treatment using Adobe Photoshop CC counting tool. A total of eight coronal SN sections were counted per animal, and the counted sections were 90 μm apart.

For stereological quantification of colorimetric IHC, unbiased count of DAB-stained neurons was performed using a Leica DM4 B microscope and Stereologer2000 software. The investigator was blind to experimental conditions. An exhaustive count of SNc TH-immunostained neurons (Slab Sampling Interval = 1, Total Number of Sections = 20, Section Sampling Interval = 2) was performed with 63X oil objective after outlining the

SNc with a 10X objective. The count was performed using a total of 100 dissectors (Frame Area: 5000 $\mu m^2$, Frame Height: 20 µm, Guard Height: 2 µm, Frame Spacing: 100 µm). A neuron was considered as positive for immunoreactivity when its nucleus fell inside the dissector borders without touching the exclusion lines. For SNr TH-immunoreactive neurons (Fig. 1E: Slab Sampling Interval = 1, Total Number of Sections = 24, Section Sampling Interval = 3; Fig. 2C: Slab Sampling Interval = 1, Total Number of Sections = 20, Section Sampling Interval = 2), a rare event protocol was used to perform an exhaustive count with a 10X objective (Frame Area: 5000 $\mu m^2$, Frame Height: 20 µm, Guard Height: 2 µm, Frame Spacing: 100 µm).

**Plasma nicotine and cotinine levels**

Blood samples were collected from adult mice after two weeks of nicotine treatment. Blood from the left ventricle was drawn from mice anesthetized with ketamine/xylazine and assayed for plasma nicotine metabolite, cotinine, titer (18 ng/mL) by high performance liquid chromatography (NMS Labs).

**Behavioral testing**

To assess PD-related behavioral deficits associated with A53T-expression and effects of nicotine treatment, mouse behavioral pattern monitor (BPM, San Diego Instruments) chambers were used to measure locomotor activity and investigatory behavior[6,7]. This system collects data encompassing total traveling distance, rearing movements, duration spent in the center, number of entries to the center, transitions (number of times mouse entered one of nine regions), and number of investigatory nosepokes (holepokes). A mouse BPM chamber is a clear Plexiglas box containing a 30 × 60 cm holeboard floor. Each chamber is enclosed in a ventilated outer box to protect it from outside ambient noise and light. The location of the mouse is obtained from a grid of 12 × 24 X-Y array of infrared photobeams that are placed 1 cm above the floor. There are 8 square sectors (15.2 cm wide) in each chamber. Crossovers between each sector are defined as movements between any of these sectors. Each chamber is also divided into 9 regions unequal in size that are used primarily to define entries into the corners and the center. Rearing is detected by an array of 16 photobeams placed 2.5 cm above the floor. Holepokes are detected by 11 1.4-cm holes in the chamber (3 in the floor and 8 in the wall), each equipped with an infrared photobeam. The status of the photobeams is sampled every 55ms. A change in the status triggers the storage of information in a binary data file together with the duration of the photobeam status. Subsequently, the raw data files are transformed into (x, y, t, event) ASCII data files composed of the (x, y) location of the animal in the mouse BPM chamber with a resolution of 1.25 cm, the duration of each event (t) and whether a holepoke or rearing occurred (event). ASCII data were then exported into Microsoft Excel files for subsequent statistical analyses with GraphPad Prism 8.4.0.

A total of eight chambers were used, each chamber measuring one mouse per session (40 min). The BPM test was conducted after 14 days of nicotine administration and performed over two days with male mice tested on the first day and female mice on the second day to avoid disruption of behavior by scent from opposite sex. The animals were brought into the testing room 1 hour before testing. During testing, a white noise generator produced background noise at 65 dB. The chambers were cleaned thoroughly between testing sessions.

For chronic nicotine exposure experiments, mice were divided into four groups (two genotypes: hα-syn+, hα-syn−; two treatments: control, nicotine). To test the effects of Nurr1 overexpression on behavior, hα-syn+ mice were divided into three groups (two genotypes: hα-syn+, hα-syn−; two types of SN viral injections: AAV.GFP, AAV.Nurr1).

**Statistical analysis**

Data were analyzed using two-tailed Student's t-test and one-way, two-way, or mixed model analysis of variance (ANOVA), as appropriate to each experiment. A criterion based on z-score was used to detect outliers prior to running the ANOVA. The level 0.01 was chosen as the decision criterion for the z-score of 3.291 beyond which a datum is considered an outlier[8]. Significant main effects and interactions were followed by Bonferroni's Multiple Comparisons test. Data are represented by mean and standard error in bar and line graphs, or by the median and interquartile range with all data points in box and whisker plot. Alpha level was set to 0.05 for all analyses. Appropriate sample size for each experiment have been determined with standard Cohens's d power analysis with target power set to 0.8 and alpha level to 0.05. Data were analyzed with IBM SPSS Statistics 26.0 and GraphPad Prism 8.4.0. Graphs were generated with Microsoft Excel and GraphPad Prism 8.4.0.

### SUPPLEMENTARY FIGURE LEGENDS

**Figure S1 – Inducible human A53T alpha synuclein accumulates over time.**

**A**, Confocal images of coronal section (30 μm) through the SN of the *PITX3-IRES2-tTA/tetO-A53T* double-transgenic mice showing a substantial increase of hα-syn expression after 90 days off doxycyline (DOX) when compared to 10 days off DOX. Colocalization shows that hα-syn was selectively expressed in the TH+ cells (arrowheads) in the substantia nigra (SN). Scale bars = 100 μm.

**B**, Quantification of neurons displaying hα-syn /TH colocalization shows a 2-fold increase in hα-syn expression after 90 days off DOX and a 3-fold increase after 120 days. The expression after 180 days off DOX was comparable to 90 days off DOX. Graphs show mean ± SEM. The number of animals is annotated in parenthesis for each condition.

**Figure S2 – Lower fluid intake does not affect body weight of nicotine-exposed mice**

**A**, Daily fluid intake per mouse over 14 days of nicotine exposure. Nicotine-treated group received 50mg/L nicotine in 1% saccharin solution, while the control group received 1% saccharin solution in place of regular drinking water.

**B**, Daily fluid intake per body weight shows that nicotine-exposed mice consumed slightly less fluid than control ($t_{(26)}$=2.61, *p<0.05).

**C**, The body weight measured at P60 and P74 indicates that 2-week nicotine exposure does not affect mouse weight.

Graphs show mean ± SEM: The number of animals is annotated in parenthesis for each condition.

**Figure S3 – SNc DAergic neurons display dendritic arborization extending onto SNr GABAergic neuropil.**

Confocal images of coronal section (30 μm) through the SN of adult (P60) VGAT-ZsGreen mouse immunolabeled with TH and NeuN markers. TH+ neurons in the SNc (pars compacta) display dendrites (arrows) extending onto the SNr and in close contact with NeuN+/VGAT+ neurons. Scale bars = 100 μm. VTA: ventral tegmental area.

**Video S1 – VGAT-ZsGreen labels VGAT+ neurons.**

Confocal images of coronal section (30 μm) through the SN of adult (P60) vesicular GABA transporter (VGAT)-ZsGreen mice were acquired with a total of 11 Z-stack layers 2 μm apart from each other. Video (40 frames/second) prepared with Imaris software (Oxford Instruments) demonstrates co-localization of VGAT (green), NeuN (red), and TH (blue) in the SNr.

### REFERENCES

1. Tamamaki, N., Yanagawa, Y., Tomioka, R., Miyazaki, J.-I., Obata, K. & Kaneko, T. Green fluorescent protein expression and colocalization with calretinin, parvalbumin, and somatostatin in the GAD67-GFP knock-in mouse. *J. Comp. Neurol.* **467**, 60–79.

2. Lin, X., Parisiadou, L., Sgobio, C., Liu, G., Yu, J., Sun, L., Shim, H., Gu, X.-L., Luo, J., Long, C.-X., Ding, J., Mateo, Y., Sullivan, P. H., Wu, L.-G., Goldstein, D. S., Lovinger, D. & Cai, H. Conditional Expression of Parkinson's Disease-Related Mutant -Synuclein in the Midbrain Dopaminergic Neurons Causes Progressive

Neurodegeneration and Degradation of Transcription Factor Nuclear Receptor Related 1. *J. Neurosci.* **32**, 9248–9264 (2012).

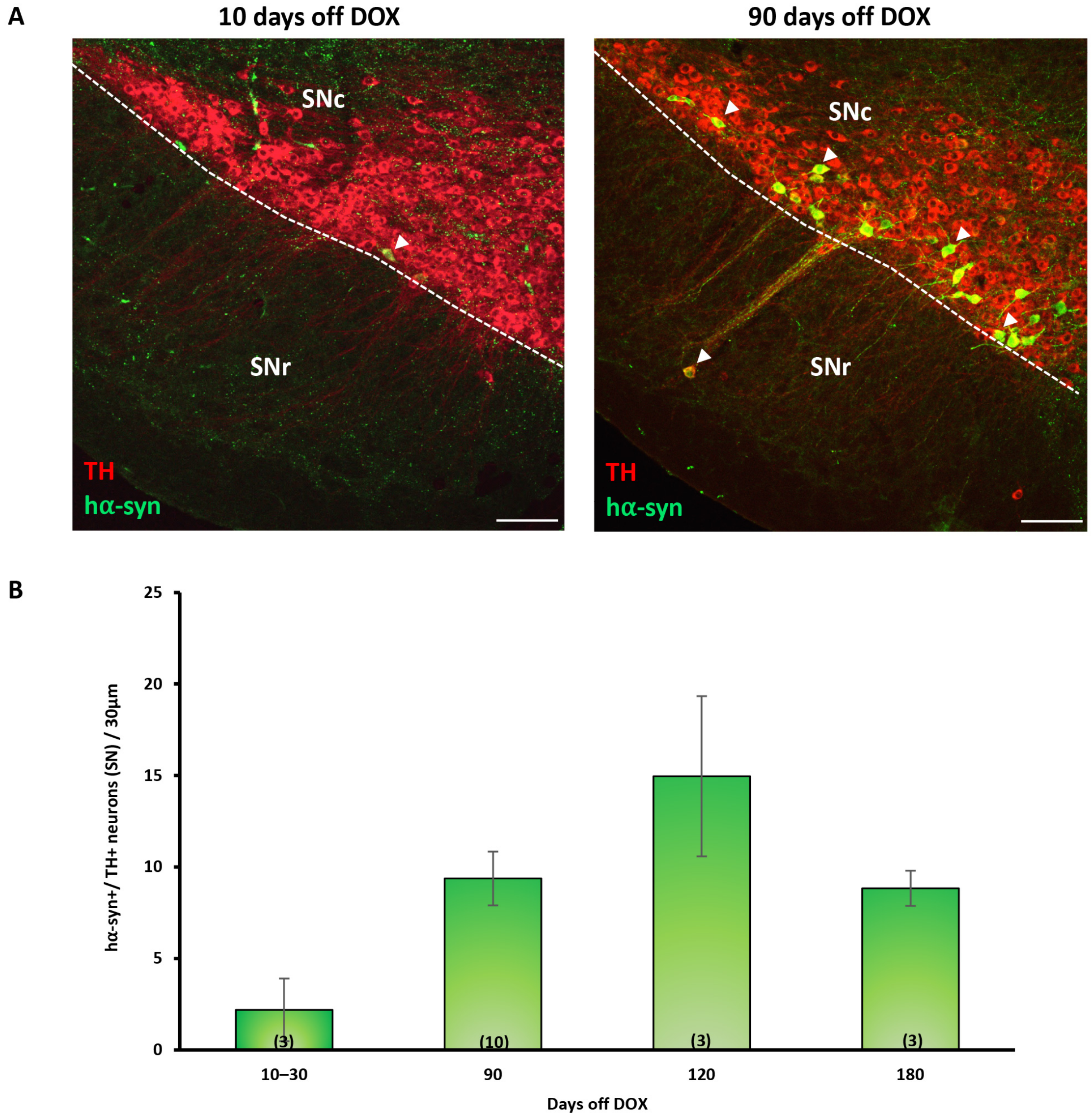
A
10 days off DOX
90 days off DOX
SNc
SNr
TH
hα-syn
B
hα-syn+/ TH+ neurons (SN) / 30µm
0
5
10
15
20
25
(3)
(10)
(3)
(3)
10–30
90
120
180
Days off DOX

**FIGURE S1**

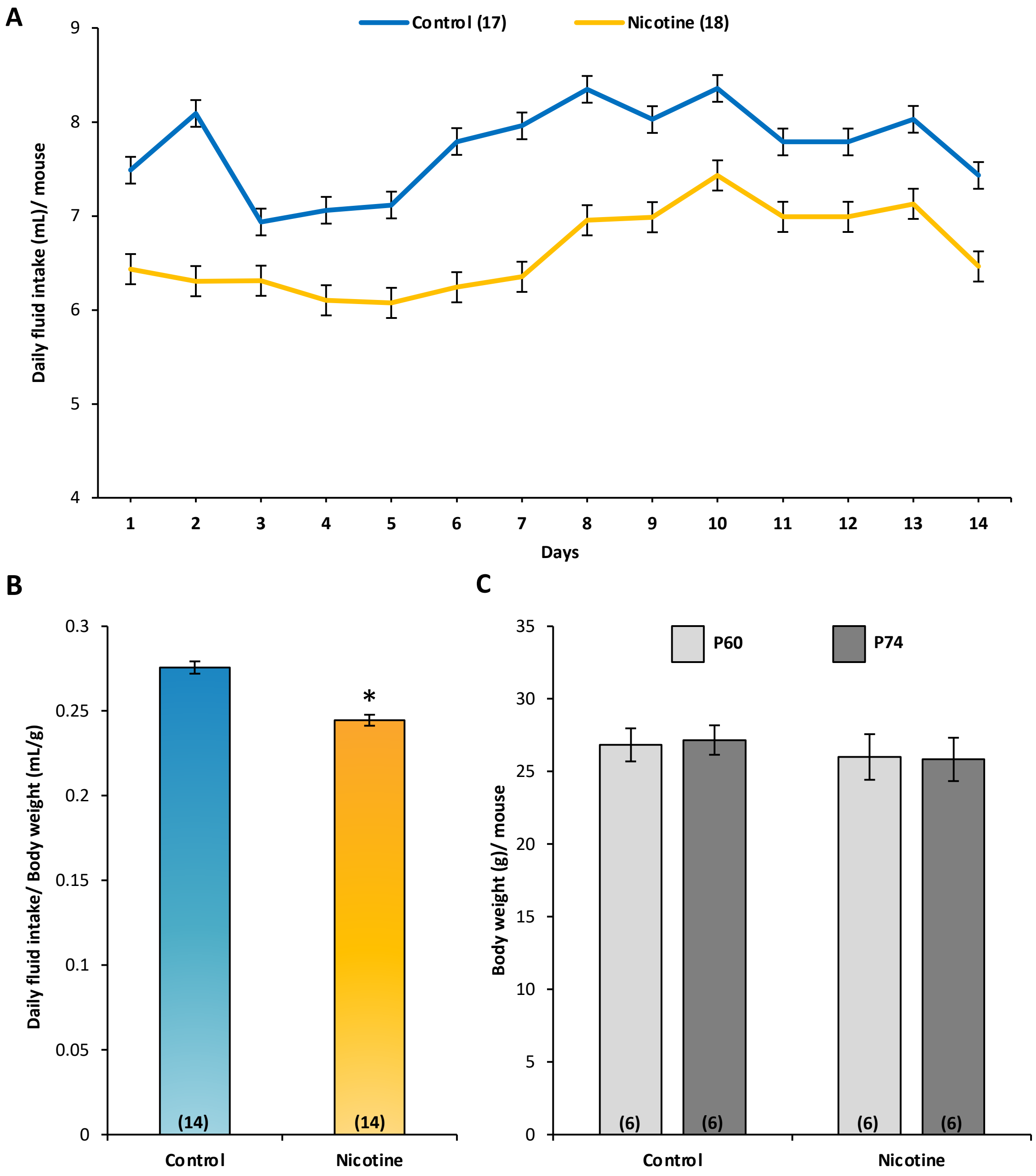
A
Control (17)
Nicotine (18)
Daily fluid intake (mL)/ mouse
9
8
7
6
5
4
1
2
3
4
5
6
7
8
9
10
11
12
13
14
Days
B
Daily fluid intake/ Body weight (mL/g)
0.3
0.25
0.2
0.15
0.1
0.05
0
*
(14)
(14)
Control
Nicotine
C
P60
P74
Body weight (g)/ mouse
35
30
25
20
15
10
5
0
(6)
(6)
(6)
(6)
Control
Nicotine

**FIGURE S2**

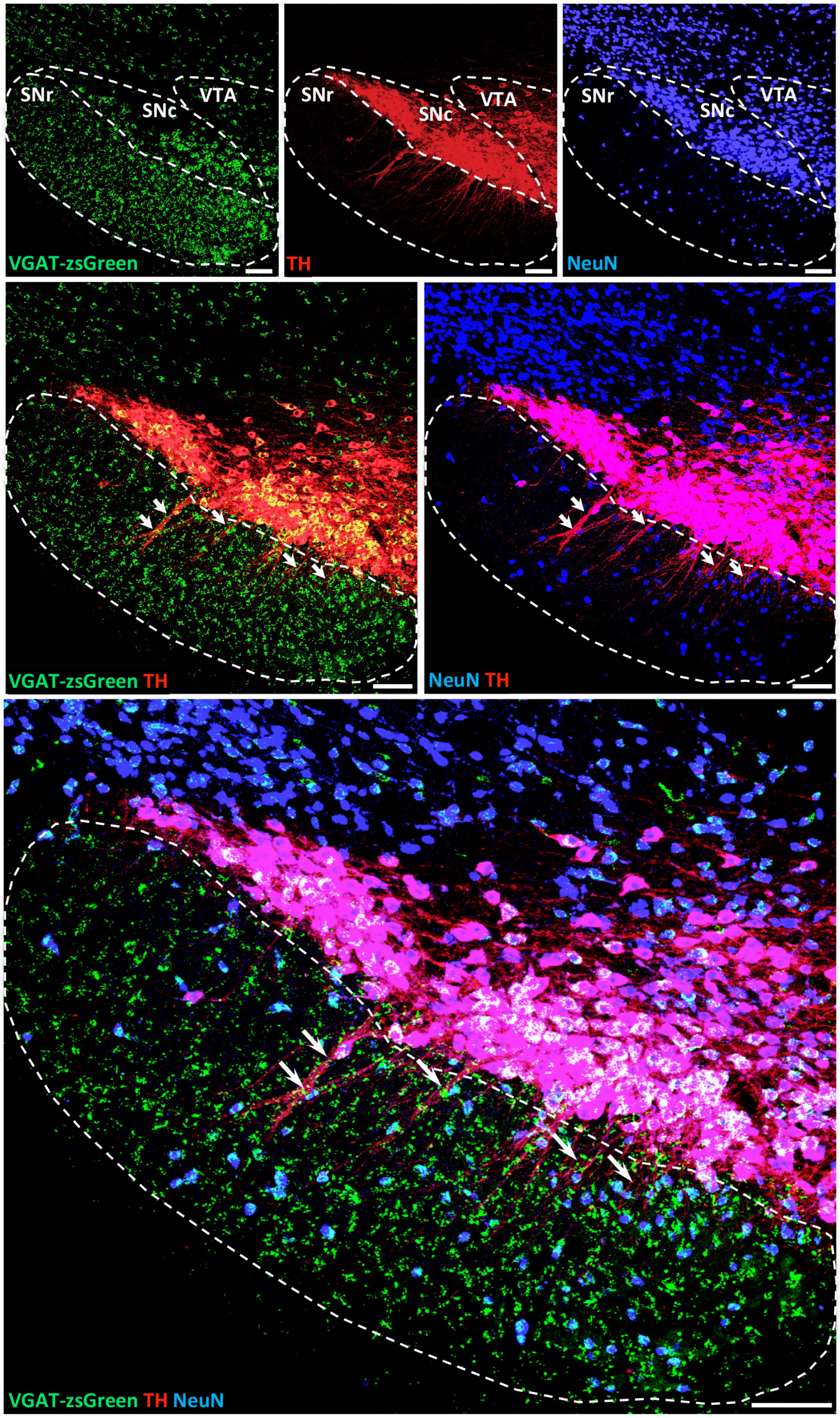
SNr
SNc
VTA
VGAT-zsGreen
SNr
SNc
VTA
TH
SNr
SNc
VTA
NeuN
VGAT-zsGreen TH
NeuN TH
VGAT-zsGreen TH NeuN

**FIGURE S3**
